# Paralogs in the PKA regulon traveled different evolutionary routes to divergent expression in budding yeast

**DOI:** 10.1101/860981

**Authors:** Benjamin Murray Heineike, Hana El-Samad

## Abstract

Functional divergence of duplicate genes, or paralogs, is an important driver of novelty in evolution. In the model yeast *Saccharomyces cerevisiae*, there are 547 paralog gene pairs that survive from an interspecies Whole Genome Hybridization (WGH) that occurred ∼100MYA. Many WGH paralogs (or ohnologs) are known to have differential expression during the yeast Environmental Stress Response (ESR), of which Protein Kinase A (PKA) is a major regulator. While investigating the transcriptional response to PKA inhibition in *S. cerevisiae,* we discovered that approximately 1/6^th^ (91) of all ohnolog pairs were differentially expressed with a striking pattern. One member of each pair tended to have low basal expression that increased upon PKA inhibition, while the other tended to have high but unchanging expression. Examination of PKA inhibition data in the pre-WGH species *K. lactis* and PKA-related stresses in other budding yeasts indicated that unchanging expression in response to PKA inhibition is likely to be the ancestral phenotype prior to duplication. Analysis of promoter sequences of orthologs of gene pairs that are differentially expressed in *S. cerevisiae* further revealed that the emergence of PKA-dependence took different evolutionary routes. In some examples, regulation by PKA and differential expression appears to have arisen following the WGH, while in others, regulation by PKA appears to have arisen in one of the two parental lineages prior to the WGH. More broadly, our results illustrate the unique opportunities presented by a WGH event for generating functional divergence by bringing together two parental lineages with separately evolved regulation into one species. We propose that functional divergence of two ohnologs can be facilitated through such regulatory divergence, which can persist even when functional differences are erased by gene conversion.

## Introduction

Gene duplication is considered an important source of novelty and adaptation in evolution (Conant and Wolfe, 2008; Des Marais and Rausher, 2008; Hittinger and Carroll, 2007; Ohno, 1970; Taylor and Raes, 2004). In particular, the duplication of entire genomes, or Whole Genome Duplications (WGD), have been hypothesized to offer a unique avenue to generate evolutionary diversity. Genes that are rarely retained as paralogs in small scale duplication events can be retained as paralogs during WGD. Paralogs resulting from a WGD are known as “ohnologs”. (Conant and Wolfe, 2007; Guan et al., 2007; van Hoek and Hogeweg, 2009; Wapinski et al., 2007).

Much of the understanding we have of WGD comes from work done on the model organism *Saccharomyces cerevisiae*, the first eukaryote to have its genome fully sequenced (Goffeau et al., 1996). Not long after the *S. cerevisiae* genome was published, researchers discovered that its lineage had undergone a WGD approximately 100Mya (Wolfe and Shields, 1997). Today, *S. cerevisiae* retains 547 ohnologs from this event, comprising over 30% of its annotated protein coding genes (Byrne and Wolfe, 2005). As more budding yeast sequences have become available, a greater understanding has developed surrounding the budding yeast WGD. One particularly surprising recent discovery was that the WGD event was most likely an allo-polyploidization, or hybridization between two species that had separated approximately 40 million prior (Marcet-Houben and Gabaldón, 2015). As a result, a more apt description for the event would perhaps be a Whole Genome Hybridization (WGH). Of the two species that hybridized during the WGH, one was closely related to the ancestor of the Zygromyces/Torulospora (ZT) branch of pre-WGH species (parent A in Fig 1A). The other likely separated from the ancestral lineage closer to the time when the ancestor of the Kluyveromyces/Lachancea/Eremothecium (KLE) branch diverged (parent B in Fig 1A).

**Figure 1:**
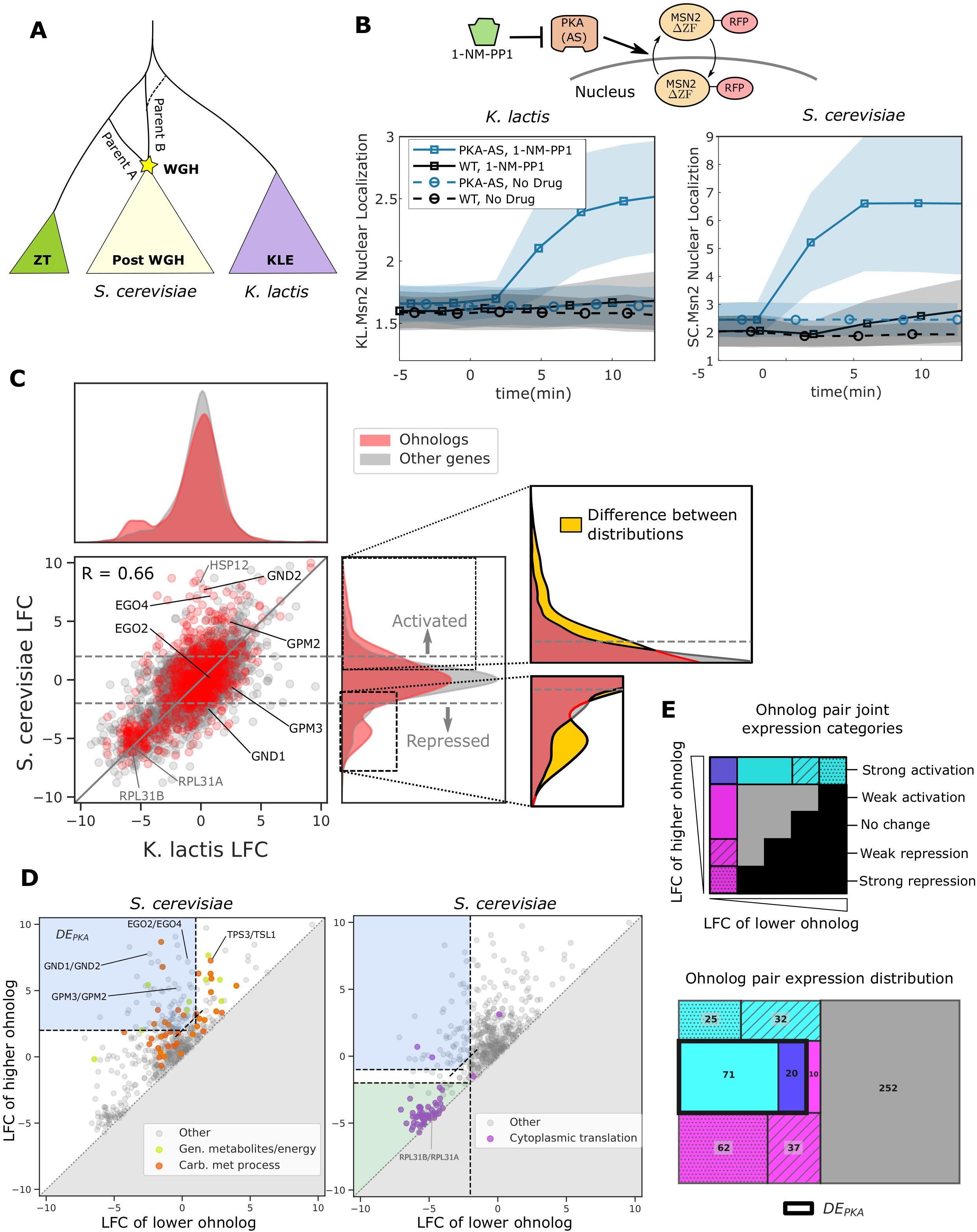
Ohnologs are enriched in genes affected by PKA inhibition in *S. cerevisiae* but in ohnolog pairs in which one member is activated, the other is generally unaffected. (A) Simplified schematic depicting the budding yeast Whole Genome Hybridization (WGH). The *Kluyveromyces/Lachancea/Eremothecium* (KLE) branch is shaded purple, and the *Zygosaccharomyces/Torulaspora* (ZT) branch is shaded green. Parent A is more closely related to the ZT branch and parent B is more closely related to the KLE branch. The dashed line illustrates the fact that the phylogenetic branch point of parent B has not been fully resolved and may be early on the ZT branch or on the KLE branch. (B) Msn2 nuclear localization in *K. lactis* (left) and *S. cerevisiae* (right) for WT and PKA Analog Sensitive (PKA-AS) strains following the addition of control media or 4uM 1-NM-PP1. Solid line represents the mean and the shaded area represents the standard deviation for at least 27 single cell measurements in *K. lactis* and at least 84 single cell measurements in *S. cerevisiae*. A diagram of the experimental system is shown. PKA-AS drives nuclear export of Msn2(ΔZF)-RFP (denoted by SC.Msn2 in *S. cerevisiae* or KL.Msn2 in *K. lactis*) and is inhibited by 1-NM-PP1. (C) Log Fold Change (LFC) comparing RNA sequencing data collected from strains in which PKA-AS was inhibited with 3uM 1-NMPP1 versus DMSO controls. Data was collected after 50 min in both *S. cerevisiae* (y-axis) and *K. lactis* (x-axis) following administration of the drug. LFC values are only shown for genes that had orthologs in both species. Genes that are ohnologs in *S. cerevisiae* are colored red, and all other genes are colored grey. The y=x line is shown in grey. Example differentially expressed ohnolog pairs highlighted later in the text are indicated with black text. The canonical Msn2/4 target HSP12, and a representative pair of ohnologs associated with cytoplasmic translation, RPL31A/B, are indicated with grey text. Horizontal dashed grey lines indicate LFC values of 2.0 and -2.0 which are thresholds used for identifying activated and repressed genes. The kernel density estimates of the distribution of LFC for ohnologs in *S. cerevisiae* and for their *K. lactis* orthologs are shown to the right and above the scatter plot respectively. The distribution of LFC for all other genes is shown in grey for comparison in both plots. The inset highlights the difference (yellow) between the LFC distribution for ohnologs and that of all other genes for genes that are either induced (upper inset) or repressed (lower inset) by PKA inhibition in *S. cerevisiae*. (D) Comparison of LFC for ohnolog pairs in *S. cerevisiae* is shown in both plots. The lower LFC value for each ohnolog pair is plotted on the x-axis and the higher LFC value is plotted on the y-axis, hence all the data lies above the y=x line. The data points are colored by selected GO-terms. The blue shading for the left plot indicates the LFC criteria for selecting differentially expressed ohnologs in which one member of the pair is induced by PKA inhibition (DE_PKA_). The blue shading for the right plot indicates LFC criteria for selecting differentially expressed ohnologs in which one member of the pair is repressed by PKA inhibition. The green shading in the right plot indicates ohnolog pairs in which both genes are repressed by PKA inhibition. (E) The top panel describes 8 categories of ohnolog pairs based on their joint LFC expression. Each axis is divided into bins based representing the level of response to PKA inhibition ranging from strong repression to strong activation. The bin for the x-axis and y-axis are chosen based on the LFC value of the lower LFC ohnolog and higher LFC ohnolog respectively. Black shading indicates combinations that do not exist because the lower LFC ohnolog always has a lower LFC value than the higher LFC ohnolog. In the treeplot (bottom panel), the area of the boxes is proportional to the number of ohnolog pairs present in each category as indicated by the color and pattern from the top panel. The thick black outline contains ohnolog pairs that belong to DE_PKA_ which includes ohnolog pairs from two sets. In the first set (containing 20 ohnolog pairs), one member is strongly activated (LFC>2.0) and the other is strongly repressed (LFC<-2.0). In the second set (containing 71 ohnolog pairs), one member is strongly activated, and the other does not change (-2.0 < LFC<1.0). This set also excludes ohnolog pairs for which the difference in expression between the high and low LFC ohnologs was less than 2.0.

Several characteristic phenotypes underwent striking changes following the WGH, including a drastic change in metabolic lifestyle between Crabtree positive yeast (post-WGH species) that use fermentation to metabolize glucose when oxygen is present and Crabtree negative yeast (most pre-WGH yeast species) that use respiration (Hagman and Piškur, 2015; Merico et al., 2007). There were also major changes in the Environmental Stress Response (ESR) following the WGH (Roy et al., 2013; Thompson et al., 2013). The ESR is a common gene expression program consisting of hundreds of genes that are either activated or repressed in a variety of stressful conditions such as glucose depletion, osmotic shock, and oxidative stress (Causton et al., 2001; Gasch et al., 2000). Previous studies have demonstrated that much of the ESR is conserved across budding yeast species, and that the level of conservation depends on whether the genes are activated or repressed in the ESR. The set of genes repressed in the ESR, which contains genes related to ribosomal biogenesis and protein production, is more conserved across different yeast lineages. By contrast, membership in the activated ESR is more variable (Roy et al., 2013; Thompson et al., 2013). Several of the evolutionary changes that occurred in the activated ESR following the WGH are consistent with changes in metabolic lifestyle between Crabtree and non-Crabtree yeast. Such changes include induction of more amino acid biosynthesis, purine biosynthesis, and oxidative phosphorylation genes as well as repression of fewer mitochondrial genes in post-WGH species compared to pre-WGH species (Thompson et al., 2013).

The Protein Kinase A (PKA) pathway is one of the primary master regulators of the ESR (Gasch et al., 2000; Mace et al., 2019; Zaman et al., 2009). Cells with a hyperactive PKA pathway, in addition to being sensitive to stress, also fail to grow on carbon sources that require respiration (Jiang et al., 1998; Toda et al., 1987). While comparative analyses of gene expression between budding yeast species have described broad patterns of conservation and divergence of the genes in the ESR that coincide with the emergence of the respiro-fermentative lifestyle (Brion et al., 2016; Roy et al., 2013; Thompson et al., 2013), the precise evolutionary rewiring of the PKA regulon itself has not been clearly delineated. Understanding the mechanisms by which this stress response pathway has been rewired could give us insight into how yeast species evolve to cope with the unique stresses of their particular ecological niches.

In this work, we probed the specific evolutionary rewiring of the PKA program following the WGH by directly inhibiting PKA in *K. lactis*, a species from a pre-WGH lineage and *S. cerevisiae*, the canonical post-WGH species, and compared the ensuing gene expression profiles. We identified 91 WGH ohnologs in *S. cerevisiae* that showed differential expression under PKA inhibition, with one ohnolog induced under this perturbation and the other with constant or even decreased expression. Furthermore, in this set, on average, the ohnolog that was induced by PKA inhibition had lower basal expression than the uninduced ohnolog. The phenotype for the shared ortholog in *K. lactis* largely resembled that of the uninduced ohnolog -- high basal expression and no induction following PKA inhibition. We explored this surprising observation in depth using publicly available gene expression data for a range of budding yeast species spanning the WGH (Roy et al., 2013; Thompson et al., 2013; Tsankov et al., 2010). Our investigations revealed that for these differentially expressed ohnologs, the ancestral transcriptional response to PKA inhibition generally featured high basal expression and low induction, motivating the need to explore the emergence of PKA dependence in the regulation of these genes.

To gain insight into the evolutionary trajectories of PKA dependence in these ohnologs, we carried out binding site analysis of their gene promoter sequences from a large set of recently published budding yeast genomes (Shen et al., 2018), focusing on the DNA binding motif for Msn2 and Msn4, the canonical transcription factors downstream of PKA. These analyses revealed that for some genes, divergence in regulation by PKA between ohnologs occurred after the WGH, while for others, differential expression arose when the ancestors of the paralogs were in different species prior to the WGH. We also identified an example in which differential regulation that arose before the WGH may have been preserved in an ohnolog pair despite a gene conversion that occurred following the WGH and homogenized the protein content. Given the abundance of allopolyploid events in diverse species from plants (del Pozo and Ramirez-Parra, 2015) to vertebrates (Matos et al., 2015) these examples represent principles for generating novelty in evolution that might be a general feature in evolution across the spectrum of life.

## Results

### A PKA analog sensitive allele in *K. lactis* allows for comparing the transcriptional response to PKA inhibition with that of *S. cerevisiae*

Information about stress and carbon source availability is transmitted to the cell through the second messenger cAMP. In favorable conditions, cAMP levels are high and the abundant cAMP binds to the regulatory subunit of the PKA hetero-tetramer, releasing the active subunit to phosphorylate PKA targets. Conversely, when cAMP levels drop, its dissociation from PKA inhibits PKA activity, allowing dephosphorylation of PKA targets. The paralogous transcription factors Msn2 and Msn4 are two prominent targets of PKA, which when dephosphorylated translocate into the nucleus and activate a majority of the genes that make up the activated portion of the ESR (Gasch et al., 2000; Görner et al., 2002; Hao and O’Shea, 2011; Stewart-Ornstein et al., 2013). Inhibition of PKA also activates the transcriptional repressors Tod6 and Dot6, which translocate into the nucleus to repress ribosomal biogenesis genes. Concurrently, the protein Sfp1, which activates ribosomal biogenesis genes, translocates out of the nucleus (Lippman and Broach, 2009). The genes regulated by Dot6, Tod6 and Sfp1 are part of the repressed ESR. As a result, inhibition of PKA in *S. cerevisiae* causes a severe slowdown of growth (Jiang et al., 1998).

To compare the response to PKA inhibition between *K. lactis,* a species that diverged prior to the WGH (pre-WGH), and the model species *S. cerevisiae* whose ancestor underwent the WGH (post-WGH), we used a gatekeeper mutation strategy to allow precise chemical control of PKA’s kinase activity (Bishop et al., 2000; Blethrow et al., 2004; Islam, 2018). For PKA in *S. cerevisiae,* this strategy consists of mutating a “gatekeeper” Methionine residue to a less bulky Glycine residue for all three PKA catalytic subunit isoforms (M164G, M147G and M165G for TPK1, TPK2, and TPK3 respectively). The resulting expanded active site still accepts ATP and allows the kinase to phosphorylate its targets with close to wild-type efficiency. However, when a bulky ATP analog (1-NM-PP1) is provided in the media, it effectively inhibits any PKA activity. This PKA analog sensitive (AS) allele strategy was previously used to investigate many properties of PKA signaling in *S. cerevisiae* (Hansen et al., 2015; Hansen and O’Shea, 2016, 2015a, 2015b; Hao and O’Shea, 2011; Mace et al., 2019; Zaman et al., 2009)

While strains with gatekeeper mutations to PKA in *S. cerevisiae* were available, there were no such strains for *K. lactis.* To inhibit PKA in *K. lactis,* we constructed a strain containing gatekeeper mutations in each of the two PKA catalytic subunit genes present in that species (M222G for KL.TPK2 and M147G for KL.TPK3). We used a single plasmid CRISPR/Cas9 gene editing strategy (Ryan and Cate, 2014), and delivered the Cas9 expression cassette and sgRNA on a universal ARS plasmid that allows for stable expression in a number of budding yeast species (Liachko and Dunham, 2014). Addition of 1-NM-PP1 to both *S. cerevisiae* and *K. lactis* strains containing PKA-AS alleles stalled growth in both species (Fig S1A) but had little effect on strains bearing the WT alleles, suggesting successful inhibition of PKA in *K. lactis* using this strategy given the known role of PKA in growth control.

To further validate our ability to inhibit PKA, we sought to measure the nuclear translocation of Msn2. The *K. lactis* ortholog of Msn2 and Msn4 (KL.Msn2) has been previously shown to be required for PKA dependent regulation of mating pathway genes (Barsoum et al., 2011). To directly investigate whether nuclear localization of KL.Msn2 is regulated by PKA as its orthologs (Msn2/4) are in *S. cerevisiae*, we integrated and constitutively overexpressed a fluorescently tagged KL.Msn2(C623S)-RFP into the genome of the PKA-AS strain. This Msn2 allele contains a mutation in a conserved cysteine in the zinc-finger domain that abolishes binding to DNA, therefore minimizing the effects of overexpressing the transcription factor (Stewart-Ornstein et al., 2013). For comparison, we also constructed a similar Msn2 reporter in a *S. cerevisiae* PKA-AS strain (SC.Msn2(C649S)-RFP). Following addition of 1-NM-PP1 to *K. lactis* PKA-AS cells, KL.Msn2-RFP became enriched in the nucleus, just as SC.Msn2-RFP did in *S. cerevisiae* (Fig 1B, Fig S1B) (Görner et al., 1998). These data indicate that the nuclear localization of Msn2, and therefore its ability to promote transcription, is regulated by PKA in both species.

### The transcriptional responses to PKA inhibition in *S. cerevisiae* and *K. lactis* share many similarities but also exhibit clear functional differences

To gain a broader understanding of the similarities and differences in the global transcriptional response to PKA inhibition between *S. cerevisiae* and *K. lactis*, we carried out mRNA sequencing for both species. We prepared sequencing libraries from exponentially growing PKA-AS cultures collected 50 minutes after providing saturating amounts of 1-NM-PP1 (3µM) or a DMSO control. We computed the Log Fold Change (LFC) in gene expression using the DESEQ2 algorithm by comparing read counts from yeast treated with drug to read counts from control cultures (Love et al., 2014). The transcriptional responses we observed were broadly correlated between the two species (Pearson Correlation = 0.66) (Fig 1C scatter plot). This was consistent with the observation that the ESR, and particularly the repressed ESR, is conserved in budding yeasts (Thompson et al., 2013).

To identify shared and species-specific functional enrichment, we first defined sets of genes that were induced or repressed by PKA inhibition in each species using a threshold that took into account both LFC and p-value (Fig S2). Focusing only on genes that had orthologs in both species, we further subdivided these target sets into subsets that were either (i) activated in both species, (ii) repressed in both species, (iii) activated in one species but not the other, or (iv) repressed in one species but not the other. We then quantified enrichment for GO-SLIM terms (*SGD Project*, 2018) for the different gene subsets using Fisher’s exact test (Table S1). For genes repressed in both species, enriched GO terms were related to protein production in the ribosome such as rRNA processing (p=2.38e-87) and cytoplasmic translation (p=4.71e-74), consistent with previous work relating PKA inhibition to a slowdown in growth (Airoldi et al., 2009; Brauer et al., 2008) and also consistent with the slowdown in growth we observed following PKA inhibition in both species (Fig S3). The genes repressed only in *K. lactis* were enriched for GO-SLIM terms related to mitochondrial translation (p=8.46e-33), in agreement with previous observations, and consistent with PKA’s involvement in the shift to respiratory metabolism in *S. cerevisiae* (Field et al., 2008; Ihmels et al., 2005; Thompson et al., 2013; Tsankov et al., 2010). The set of genes induced in *K. lactis* but not in *S. cerevisiae* was enriched for meiotic cell cycle control (p=2.19e-3) and conjugation (p=5.86e-3) consistent with the notion that *K. lactis* incorporates nutritional signals into its decision to undergo meiosis (Booth et al., 2010). The set of genes induced by PKA inhibition in both species was enriched for carbohydrate metabolic process (p=2.89e-6), oligosaccharide metabolic process (p=4.66e-4), and vacuole organization (including a number of autophagy related genes) (p=1.47e-3). Enrichment for these terms indicates that PKA inhibition triggers a set of conserved gene expression changes related to metabolic processes. The set of genes that were specifically induced in *S. cerevisiae* but not in *K. lactis* following PKA inhibition was also enriched for carbohydrate metabolic processes (p=8.56e-6), but in addition included a significant enrichment for genes involved in generation of precursor metabolites and energy (p=4.71e-10), response to oxidative stress (1.73e-6), and cellular respiration (2.21e-5). This is again consistent with the role of PKA inhibition in initiating respiratory metabolism in *S. cerevisiae*.

### Many ohnolog pairs are differentially activated by PKA inhibition in *S. cerevisiae*

Since *S. cerevisiae* has undergone a WGH, we were curious whether its surviving ohnologs were equally represented in the PKA regulon compared to other genes that didn’t maintain their paralogous copy from the WGH. Interestingly, we found that ohnologs were enriched both in the set of genes that were activated by PKA inhibition (p=2.67e-10) and those that were repressed by PKA inhibition (p=4.88e-6), but not in the set of genes whose expression didn’t change during PKA inhibition (Table S2, Fig 1C insets).

To further explore this observation and determine whether both members of the *S. cerevisiae* ohnolog pairs that showed enrichment exhibited the same behavior, we plotted the LFC of ohnolog pairs against one another, ordered by their LFC values (Fig 1D). To find ohnolog pairs with differential expression, we first identified sets of ohnolog pairs where one member of the pair was either activated or repressed more than four-fold (LFC > 2.0 or LFC<-2.0, respectively) with a log10(pvalue) less than -1.5. We then retained from these sets only ohnolog pairs in which the other member of the pair did not change in the same direction. Specifically, for the set of ohnolog pairs that had one member activated, we required the other member to have an LFC less than 1.5, and for pairs that had one member repressed, we required the LFC for the other member to be greater than -1.5. We further required that the difference of LFC between the two ohnologs be greater than 2.0 (Fig 1D, Table S3). Of the 509 ohnolog pairs in the *S. cerevisiae* genome which were present in our dataset, 129 ohnolog pairs had at least one ohnolog repressed and 148 had at least one ohnolog activated according to these criteria (Fig 1E). Of the 129 ohnolog pairs that had at least one ohnolog represssed, only 30 pairs (23%) were differentially expressed, with one member of the pair being repressed and the other remaining unchanged or slightly activated. By contrast, of the 148 ohnolog pairs that had at least one ohnolog activated, 91 of them (61%) were differentially expressed in response to PKA inhibition. Of that group of 91 differentially expressed ohnolog pairs, 20 had one member activated and the other repressed, while the rest had one member activated and the other unchanged. These data demonstrate that ohnolog pairs with at least one member activated by PKA inhibition were more likely to have divergent responses than those with at least one member repressed by PKA inhibition (Fig 1E).

Functional examination of the different ohnolog pairs showed that those in which both ohnologs were repressed by PKA inhibition were enriched for ribosomal genes. Of the 129 ohnolog pairs that had at least one member repressed, 61 pairs had both members repressed and all but 10 of them had at least one member associated with the GO-slim term “cytoplasmic translation” (Fig 1D right panel). By contrast, only 2 of the other 68 ohnolog pairs with one member repressed were associated with this term. These ribosomal ohnolog pairs presumably make up much of the signal that was observed in previous work describing enrichment for ribosomal proteins in retained *S. cerevisiae* ohnologs (Blanc and Wolfe, 2004; Papp et al., 2003; Seoighe and Wolfe, 1999).

Another prominent difference between the ohnologs repressed by PKA inhibition and those activated by PKA inhibition was in whether their orthologs in *K.lactis* responded similarly to PKA inhibition. Considering only ohnolog pairs that had a *K.lactis* ortholog and for which we had expression data in our dataset, there were 122 ohnolog pairs in which one member was repressed by PKA inhibition in *S. cerevisiae*. Of those, 77 (63%) had a *K. lactis* ortholog that was also repressed. By contrast, of the 138 ohnolog pairs in which one member was activated by PKA inhibition in *S. cerevisiae*, only 28 (20%) had *K. lactis* orthologs that were also activated.

Because of the intriguing pattern in the 91 differentially activated ohnolog pairs (which we will simply refer to as DE_PKA_ in the following), we turned our focus to exploring them exclusively in more depth. First, we probed their basal expression because lack of activation for one member of each ohnolog pair in this DE_PKA_ set could correspond to either low or high basal expression. To determine which scenario was more applicable, we examined basal expression of DE_PKA_ genes with and without PKA inhibition, which we quantified using average regularized log counts (rlog) data (Love et al., 2014). In the activated members of each of the DE_PKA_ ohnolog pairs, which we refer to as the DE_PKA_ high-LFC ohnologs, the basal expression had a distribution with a noticeably lower mean than that of all genes in the genome (4.3 vs. 5.9 rlog) (Fig 2A left panel, Fig S4 middle column). Furthermore, upon PKA inhibition, the distribution of expression of this group shifted clearly higher than that of all other genes (mean = 7.3 rlog), recapitulating the induction for which they were defined (Fig 2A left panel, Fig 2B left column). By contrast, the distribution of basal expression values for the DE_PKA_ genes that were unchanged or repressed following PKA inhibition (DE_PKA_ low-LFC ohnologs) was higher (mean = 6.9 rlog) and shifted down upon PKA inhibition (mean = 6.1 rlog) (Fig 2A middle panel, Figs 2B and S4 middle column). Importantly, the distribution of basal expression of the DE_PKA_ high-LFC ohnologs had a clear lower median than that of the DE_PKA_ low-LFC ohnologs (Fig 2C top panel). This indicates that the difference in PKA responsiveness between these two sets is the result of low basal expression and strong induction for the DE_PKA_ high-LFC ohnologs, and high basal expression that was either slightly reduced or unchanged by PKA inhibition for DE_PKA_ low-LFC ohnologs. *K. lactis* contains orthologs for 87 of the 91 ohnolog pairs in the DE_PKA_ set. As a group, these orthologs had a similar phenotype to that of the DE_PKA_ low-LFC ohnologs, displaying high basal expression (Fig 2A right panel, 2C lower panel) and little change in expression under PKA inhibition (Fig 2A right panel, 2B right column).

**Figure 2:**
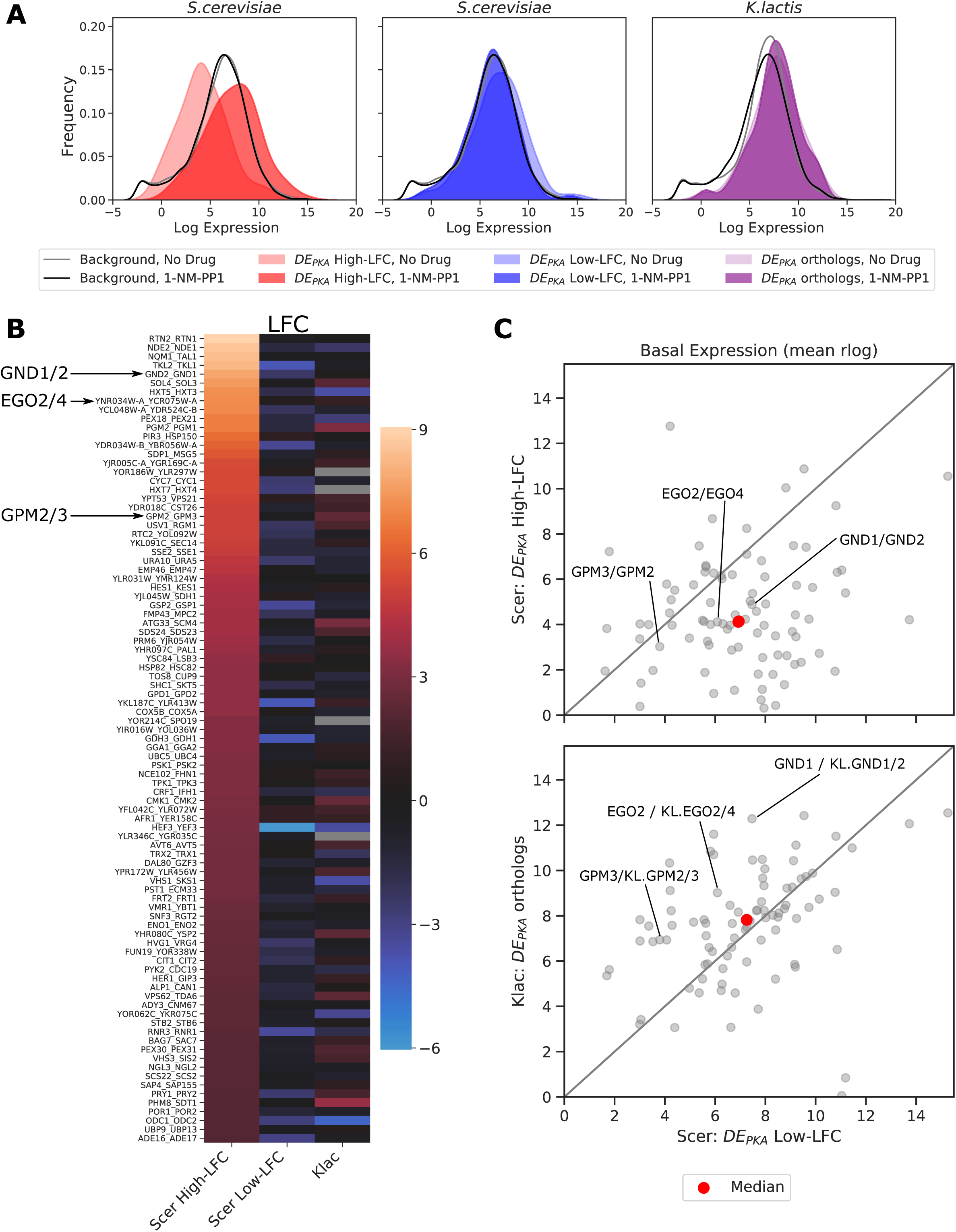
In *S. cerevisiae,* the high-LFC ohnologs from DE_PKA_ tend to have lower average basal expression while the low-LFC ohnologs tend to have higher average basal expression. Their shared orthologs from *K. lactis* on average resemble the DE_PKA_ low-LFC ohnologs. (A) Kernel density estimates for average rlog data. DE_PKA_ high and low LFC ohnologs are plotted without (light shading) and with (dark shading) treatment with 3µM 1-NM-PP1. Background expression without and with 3µM 1-NM-PP1 is also shown (grey and black lines, respectively). The background set for *S. cerevisiae* includes all annotated genes except dubious orfs identified in SGD. The background set for *K. lactis* includes all annotated genes with orthologs in *S. cerevisiae*. (B) LFC for high-LFC ohnologs (first column) and low-LFC ohnologs (middle column) from the DE_PKA_ ohnolog set in *S. cerevisiae* as well as their *K. lactis* orthologs (last column). Rows are sorted from highest to lowest LFC for the high-LFC ohnolog. Example ohnolog pairs referred to later in the text are highlighted. (C) Plot of basal expression (rlog normalized estimate of counts for no-drug samples) for DE_PKA_ low-LFC ohnologs in *S. cerevisiae* (x-axis) vs. DE_PKA_ high-LFC ohnologs (y-axis, top panel) and their orthologs in *K. lactis* (y-axis, bottom panel). Median values are shown in red, and the y=x line is shown in grey.

Taken together, our data and analyses so far reveal a class of ohnologs, amounting to about 1/6^th^ of all the retained ohnolog pairs, or about 3% of all genes in *S. cerevisiae*, which show differential expression under PKA inhibition with one member of an ohnolog pair exhibiting high induction and the other showing either no change or repression. On average, the PKA insensitive ohnolog exhibits high basal expression. The *K. lactis* orthologs of these differentially expressed ohnologs have a similar lack of induction and high basal expression. This observation posed the hypothesis that the ancestral phenotype for many of these ohnolog pairs might have been one with high basal expression and no regulation by PKA.

### Analysis of stress-response data across species suggests that high basal expression was the ancestral state for most DE_PKA_ genes

Without gene expression measurements following PKA inhibition in a wider phylogenetic range of extant species, testing whether low LFC in response to PKA inhibition was the ancestral state of the DE_PKA_ genes cannot be done directly. However, we can explore it meaningfully across a range of 15 budding yeast species spanning the WGH (Fig 3A) by capitalizing on available gene expression datasets collected in response to growth to saturation and various stress conditions known to repress the PKA pathway in *S. cerevisiae* (Roy et al., 2013; Thompson et al., 2013). To extract the conditions that most resemble PKA inhibition, we compared expression changes in *S. cerevisiae* and *K. lactis* in these datasets to our PKA inhibition data. This comparison identified five conditions (heat shock at 30 and 45 minutes, Diauxic Shift, Post Diauxic Shift, and Plateau) that were strongly correlated with PKA inhibition in both species (Pearson correlation R > 0.65 with *K. lactis* PKA inhibition and R> 0.75 with *S. cerevisiae* PKA inhibition) (Fig S5). We adopted the LFC reported for these 5 conditions (normalized per Materials and Methods) as surrogates for PKA inhibition across the species present in this dataset.

**Figure 3:**
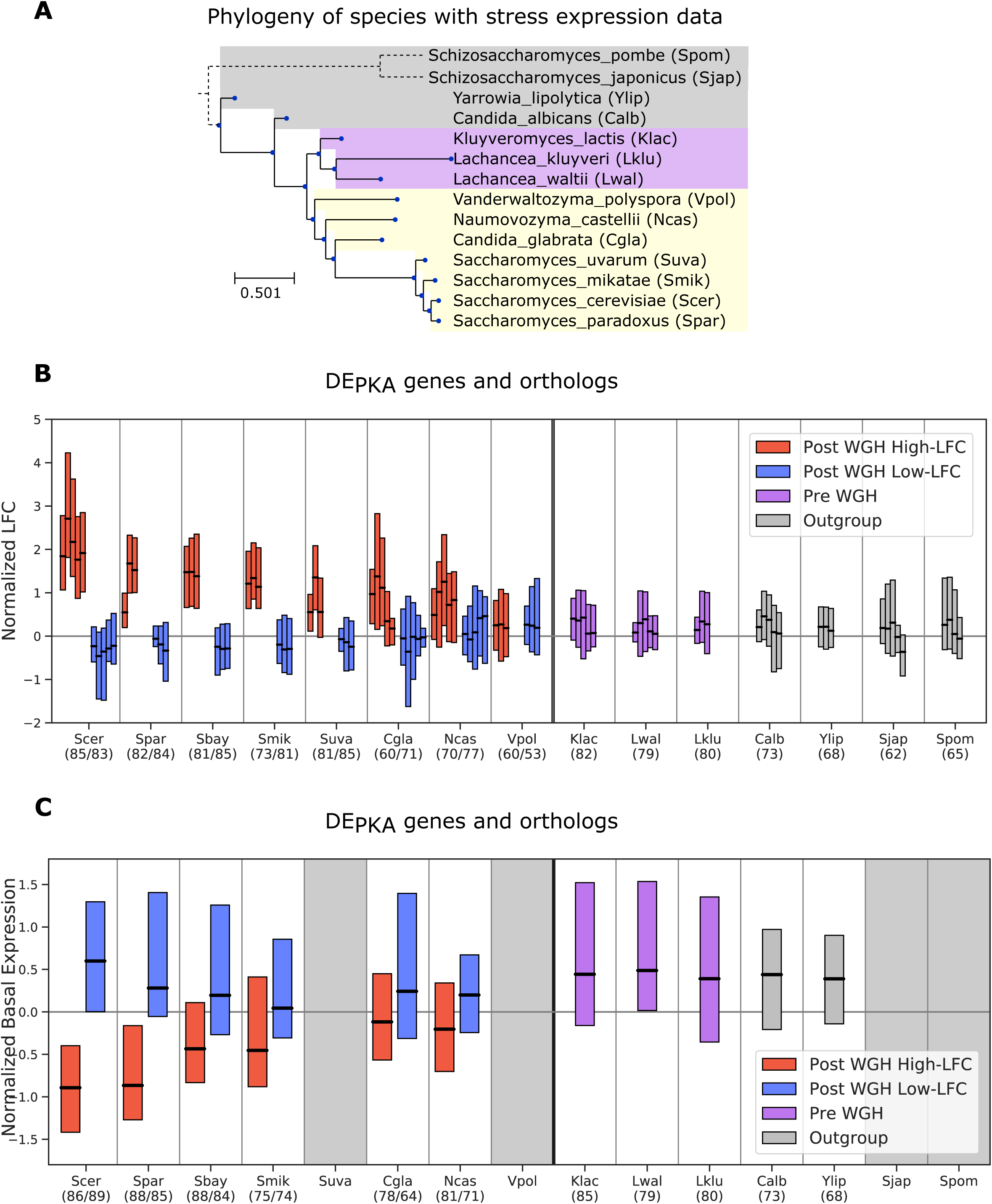
For DE_PKA_ genes, low LFC in response to PKA related stress and high basal expression is the ancestral phenotype. (A) Phylogeny of species with stress expression data used in panel (B) using the time calibrated tree generated in (Shen et al., 2018). *S. bayanus* is not shown because it is a hybrid between *S. eubayanus* and *S. uvarum* with some genes from *S. cerevisiae* (Libkind et al., 2011). *S. pombe* and *S. japonicus* branches are not drawn to scale. Boxplots in panels (B) and (C) show median Q1-Q3 range for the datasets described below for DE_PKA_ genes in *S. cerevisiae* and their orthologs in each indicated species (when present). Blue and red bars indicate low-LFC and high-LFC ohnologs (respectively) and their syntenic orthologs in Post-WGH species. Purple and grey bars are for the shared orthologs in Pre-WGH *Saccharomycetaceae* species and outgroups respectively. Numbers in parentheses indicate the number of retained orthologs. Syntenic ortholog assignment for Post-WGH species is based on the YGOB database (Byrne and Wolfe, 2005). Boxplots in panel (B) show normalized gene expression (see Materials and Methods) for the conditions most closely related to PKA inhibition in *S. cerevisiae* and *K. lactis* from (Roy et al., 2013; Thompson et al., 2013) (Fig S5). Where there are three bars, the conditions are ‘DS/LOG’, ‘PS/LOG’, and ‘PLAT/LOG’ from (Thompson et al., 2013) and where there are five bars, the conditions are those three conditions plus ‘heat shock_030’ and ‘heat shock_045’ from (Roy et al., 2013). *S. pombii* was missing ‘heat shock_045’. Boxplots in panel (C) show normalized raw expression data (see Materials and Methods) from microarray experiments comparing mRNA under exponential growth conditions to genomic DNA from (Tsankov et al., 2010). Grey regions indicate species for which no data exist. Species abbreviations: Scer = *Saccharomyces cerevisiae*, Spar = *Saccharomyces paradoxus*, Sbay = *Saccharomyces bayanus,* Smik = *Saccharomyces mikatae*, Suva = *Saccharomyces uvarum*, Cgla = *Candida glabrata*, Ncas = *Nakaseomyces castellii,* Vpol = *Vanderwaltozyma polyspora*, Klac = *Kluyveromyces lactis*, Lwal = *Lachancea waltii*, Lklu = *Lachancea kluyveri*, Calb = *Candida albicans*, Ylip = *Yarrowia lipolytica*, Sjap = *Schizosaccharomyces japonicus*, Spom = *Schizosaccharomyces pombe*.

We curated the orthologs of the 91 DE_PKA_ ohnolog pairs in the various species and compared their gene expression values in the PKA-related stress conditions. The ohnologs pairs of the DE_PKA_ set generally showed the same phenotypes in *S. cerevisiae* under these stress conditions as in our PKA inhibition dataset—high LFC for one member of the pair and low LFC in the other (Fig 3B). This is expected given our criteria for selecting these stress conditions. In the post-WGH species closely related to *S. cerevisiae*, the phenotype was also conserved. Orthologs with corresponding genomic context (or syntenic orthologs) for the high-LFC ohnologs had higher LFC, and syntenic orthologs of the low-LFC ohnologs had lower LFC on average. The separation in LFC between the syntenic orthologs of the high-LFC and low-LFC ohnologs declined in more distantly related post-WGH species. In *V. polyspora,* the most distantly related post-WGH species, the orthologs of the low and high LFC ohnologs both had low LFC in stress conditions (Fig 3B, Fig S6), although confident assignment of syntenic orthologs becomes difficult at that evolutionary distance. Importantly, for pre-WGH species, the shared orthologs of the DE_PKA_ ohnolog pairs have low LFC under these PKA-related stress conditions (Fig 3B), consistent with our hypothesis that low LFC is generally the ancestral response to PKA inhibition for DE_PKA_ ohnolog pairs.

To explore basal expression of orthologs of the DE_PKA_ genes in these species, we turned to a different dataset that measured gene expression during exponential growth on rich media using microarray data for a range of budding yeasts (Tsankov et al., 2010). After appropriate normalization (see Materials and Methods), we checked concordance of these data to rlog estimates from our RNA seq count data for *S. cerevisiae* and *K. lactis* strains under control conditions. We observed high correlation despite the different methods by which gene expression data was collected (Pearson Correlation R=0.69 and 0.60 for *S. cerevisiae* and *K. lactis* respectively) (Fig S7A). Reassuringly, there was high correlation between both datasets for the DE_PKA_ genes (R=0.67 for the low-LFC ohnologs and 0.70 for the high-LFC ohnologs) as well as for their orthologs in *K. lactis* (R=0.59) (Fig S7B), instilling confidence that the various sets are coherent in their content and can be used for cross-comparison.

With these data in hand, we could compare basal expression of DE_PKA_ genes and their syntenic orthologs across species. For the post-WGH species, the syntenic orthologs of the low-LFC DE_PKA_ ohnologs generally had a higher basal expression while those of the high-LFC DE_PKA_ ohnologs generally had lower basal expression for species more closely related to *S. cerevisiae* and higher basal expression for species more distantly related. For the pre-WGH species, the basal expression of the shared ortholog of a given DE_PKA_ ohnolog pair was typically high, similar to that of the low-LFC DE_PKA_ ohnolog in *S. cerevisiae* (Figs 3C, S8). These results were robust to redefining the ohnolog pairs of interest based on the PKA-related stress conditions rather than on the PKA-inhibition dataset (Fig S9, Figs S10 and S11 top panel). This evidence supports the notion that the basal expression level of the shared ancestors of the DE_PKA_ ohnolog pairs was, on average, high.

The use of stress conditions also allowed us to define differentially expressed ohnolog pairs from the perspective of post-WGH species other than *S. cerevisiae*. This enabled us to test whether our conclusions about the state of the ancestral phenotype were sensitive to the species used as a reference. We therefore defined sets of differentially expressed ohnologs in *N. castellii* and *V. polyspora* using gene expression for the five PKA-related conditions (see Materials and Methods for details). As when *S. cerevisiae* was used as a reference, orthologs in pre-WGH species of these ohnologs also had low LFC values (for ohnologs defined using *N. castellii* and *V. polyspora*) and high basal expression (for ohnologs defined using *N. castellii*) (Figs S10, S11).

Taken together, these data strongly support the notion that the ancestral state of ohnologs that are differentially expressed in response to PKA was characterized by high basal expression and insensitivity to PKA. The derived phenotype would therefore be characterized by high LFC in response to PKA inhibition and low basal expression. These data, however, do not indicate whether this derived phenotype for any particular ohnolog arose before the WGH in the lineage leading to the ZT branch, or after the WGH. That is because the pre-WGH species for which stress response data was present all diverged from the *S. cerevisiae* lineage at the same time or earlier than *K. lactis*. To be made with certainty, this inference would require knowledge of gene expression in the *Zygromyces/Torulospora* (ZT) branch in response to PKA related stresses. Since no such data exist, we next turned to exploring whether the promoters of the DE_PKA_ genes have a bioinformatic signal that might correlate with these phenotypes, capitalizing on the wealth of sequenced genomes in the budding yeast subphylum to formulate hypotheses about their evolutionary trajectory (Shen et al., 2018).

### The STRE is enriched in the promoters of the genes induced by PKA inhibition in *S. cerevisiae,* and *K. lactis*

To generate a framework for evaluating the characteristics of DE_PKA_ gene promoters, we first investigated bioinformatic signals associated with the promoters of all genes activated by PKA inhibition. Comparing the promoters of all genes activated under PKA inhibition in *S. cerevisiae* against the promoters of all *S. cerevisiae* genes, we identified the Stress Response Element (STRE, CCCCT), the canonical binding sequence for Msn2 and Msn4, as the most heavily enriched motif (E-value 3.3e-40) (Fig 4A) (for more detail on the promoter analysis see Supplementary Materials). A search for exact matches of the STRE binding site confirmed that, relative to all promoters in *S. cerevisiae*, PKA targets in *S. cerevisiae* were enriched for promoters that contained one or more STREs (75.2% vs. 43.9%, p-value 1.5e-16). They also had a notable increase in the average number of STREs per promoter (1.32 vs. 0.62) (Fig 4B, left). This is consistent with previous findings, implicating Msn2 and Msn4 as prominent targets downstream of PKA in *S. cerevisiae* (Görner et al., 2002; Smith, 1998). The promoters of PKA targets were also enriched for the TATA box (TATA(A/T)A(A/T)(A/G)) (70.1% with 1 or more TATA box vs. 57.5% in the promoters of all genes, p-value 2.8e-3) (Fig 4A, S13A).

**Figure 4:**
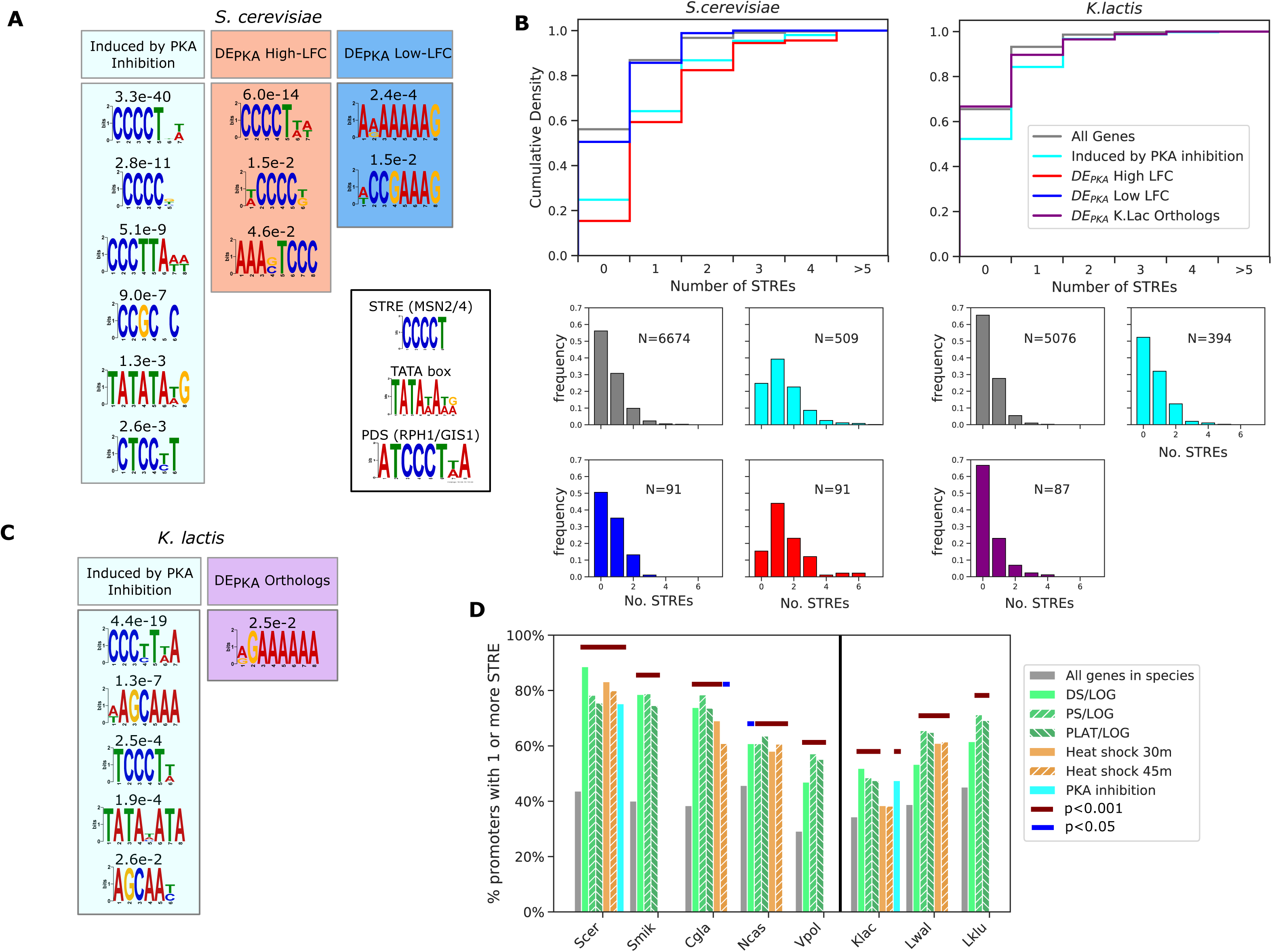
The STRE is enriched in the promoters of DE_PKA_ high-LFC ohnologs, as well as in the promoters of genes induced by PKA inhibition and PKA-related stress in various species. (A) Motifs identified using the DREME algorithm from the MEME suite (Bailey, 2011) as enriched in the promoters of all *S. cerevisiae* genes activated by PKA inhibition (light blue box) as well as DE_PKA_ high-LFC ohnologs (red box) and low-LFC ohnologs (blue box) versus a background of all *S. cerevisiae* genes. We define promoters as 700bp upstream of the start codon for this analysis. E-values are indicated above each motif. The inset shows motifs for the STRE, TATA box, and PDS. (B) Cumulative distributions (top) and histograms (bottom) of the numbers of STREs in the promoters of all genes activated by PKA inhibition as well as DE_PKA_ high and low-LFC ohnolog sets (for *S. cerevisiae,* left) or their orthologs (for *K. lactis*, right). (C) Motifs identified using the DREME algorithm from the MEME suite in promoters of all *K. lactis* genes activated by PKA inhibition (light blue box) as well as the orthologs of the DE_PKA_ ohnolog pairs (purple box) versus a background of all *K. lactis* genes. (D) Percentage of promoters with 1 or more STRE for all genes in a given species, or for genes that have LFC values above 2.5 for stresses related to PKA inhibition (data from (Roy et al., 2013; Thompson et al., 2013)). Black line separates pre-WGH from post-WGH species. Thick lines above the bars mark conditions for which number of STREs greater than 1 was statistically different than all genes in that species using Fisher’s exact test. Magenta: p-value<1.0e-3, blue: p-value<5.0e-2.

Probing the bioinformatic characteristics of promoters of genes that are responsive to PKA inhibition in *K. lactis* revealed that those also had more STREs on average compared to the promoters of all genes in that species. However, the strength of the signal was lower than in *S. cerevisiae* (E-value of 4.4e-19 vs. 3.3e-40), and the top motif we identified resembled a Post Diauxic Shift (PDS) motif (ATCCCT(T/A)A) (Pedruzzi et al., 2000) as well as an STRE (Fig 4C). Furthermore, a search for exact matches to the STRE in PKA targets in *K. lactis* revealed a weaker enrichment (47.7% of promoters with 1 or more STRE in the promoter in PKA activated genes vs. 34.5% in all genes, p=1.6e-4) (Figs4B right) than in *S. cerevisiae.* In *K. lactis*, as in *S. cerevisiae*, the promoters of genes activated by PKA were enriched for TATA boxes (70.8% with 1 or more TATA box vs. 54.1% in all genes, p=3.8e-4) (Fig S13B).

Taken together, these results indicate that the presence of STRE and TATA box motifs in promoters might be a useful proxy for activation by PKA inhibition across budding yeast species. We therefore looked for these features in the promoters of DE_PKA_ ohnologs and their *K. lactis* orthologs to see whether the presence of STRE and TATA box motifs correlated with the response to PKA we observed in our experiments. The distributions for the number and location of STREs for the DE_PKA_ high-LFC ohnologs closely resembled those of the promoters for genes activated by PKA inhibition. However, the STRE distribution for the DE_PKA_ low-LFC ohnologs resembled that of promoters of all *S. cerevisiae* genes (Figs 4B left, S12A), consistent with the fact that they were not activated in response to PKA inhibition. Furthermore, the distribution of the number of STREs in the promoters of *K. lactis* orthologs of the DE_PKA_ genes was close to that of all genes in *K. lactis* (33% vs. 34.5% had one or more STRE in the promoter), again, consistent with the fact that these genes had low LFC in response to PKA inhibition (Fig 4B right).

The number of TATA-boxes in the promoters of the DE_PKA_ high-LFC ohnologs were also increased relative to those of the promoters of all genes. However, unlike for the STRE, this enrichment was also present for DE_PKA_ low-LFC ohnologs and the *K. lactis* orthologs of the DE_PKA_ genes (Fig S13). Based on that observation, we reasoned that, at least in the context of the DE_PKA_ genes and their orthologs, the TATA box was not linked strongly enough to induction following PKA inhibition and was likely to be an ambiguous evolutionary signal. Therefore, we focused instead on the presence of STREs as a bioinformatic proxy for gene induction in response to PKA inhibition.

### Analysis of STREs in the promoters of DE_PKA_ orthologs across species revealed enrichment of STREs in the ZT branch

To investigate whether the relationship between activation by PKA inhibition and STRE presence in the promoter holds in other budding yeast species, we revisited the cross-species gene expression datasets (Roy et al., 2013; Thompson et al., 2013) and scored STRE enrichment in the promoters of genes activated under conditions correlated with PKA inhibition in *S. cerevisiae* and *K. lactis* (Fig 4D). With the exception of heat stress in *K. lactis* and diauxic shift in *L. waltii* and *L. kluyverii*, the promoters of the genes activated under these stresses had more STREs than all genes in that species (p-value<0.05 using Fisher’s Exact test) although the overall background level of STREs and the level of enrichment varied widely between species. These results suggest that there is a relationship between the presence of STREs in the promoter and gene induction under PKA inhibition that can be detected bioinformatically across budding yeast that.

We therefore quantified STREs in the promoters for orthologs of DE_PKA_ genes to search for examples in which the STRE conservation patterns might provide some insight into the evolution of gene responsiveness under conditions of PKA inhibition. We analyzed promoters in 32 pre-WGH and 12 post WGH species, including species for which no stress or PKA inhibition gene expression data exists (e.g. species in the ZT branch). For this analysis, we only focused on the DE_PKA_ ohnolog pairs in *S. cerevisiae* that had at least one STRE in the promoter of the high-LFC member. We discarded genes with high-LFC but no STRE in the promoter, reasoning that they are induced through a mechanism that doesn’t require the STRE (for example, not via Msn2/4), and thus the conservation of the STRE would not be the most relevant bioinformatic signal. We also removed pairs that had more than one ortholog in more than 8 pre WGH species which included ohnolog pairs that were present as duplicates before the WGH (such as the hexose transporters) and genes that may have undergone a separate ancient duplication which would complicate our analysis (such as SNF3/RGT2). Finally, we removed pairs that had no orthologs in pre-WGH species (such as USV1/RGM1) (see Materials and Methods).

This amounted to a total of 60 remaining ohnolog pairs. Exploration of the promoters of the orthologs of the high-LFC ohnologs in this set showed an enrichment of STREs in post WGH species (Fig 5 barplot, red bars). This enrichment was not present in the low-LFC ohnologs (Fig 5 barplot blue bars). This pattern was more prominent for species closely related to *S. cerevisiae*, with increased variability for species more distantly related and no enrichment for *V. polyspora*, the most distant post-WGH species we analyzed. *C. glabrata* was an exception to this pattern with enrichment for STREs in both the syntenic orthologs of the low-LFC and high-LFC ohnologs. This pattern is consistent with several STREs arising in the promoter of the ancestor of the high-LFC ohnolog following the WGH.

**Figure 5:**
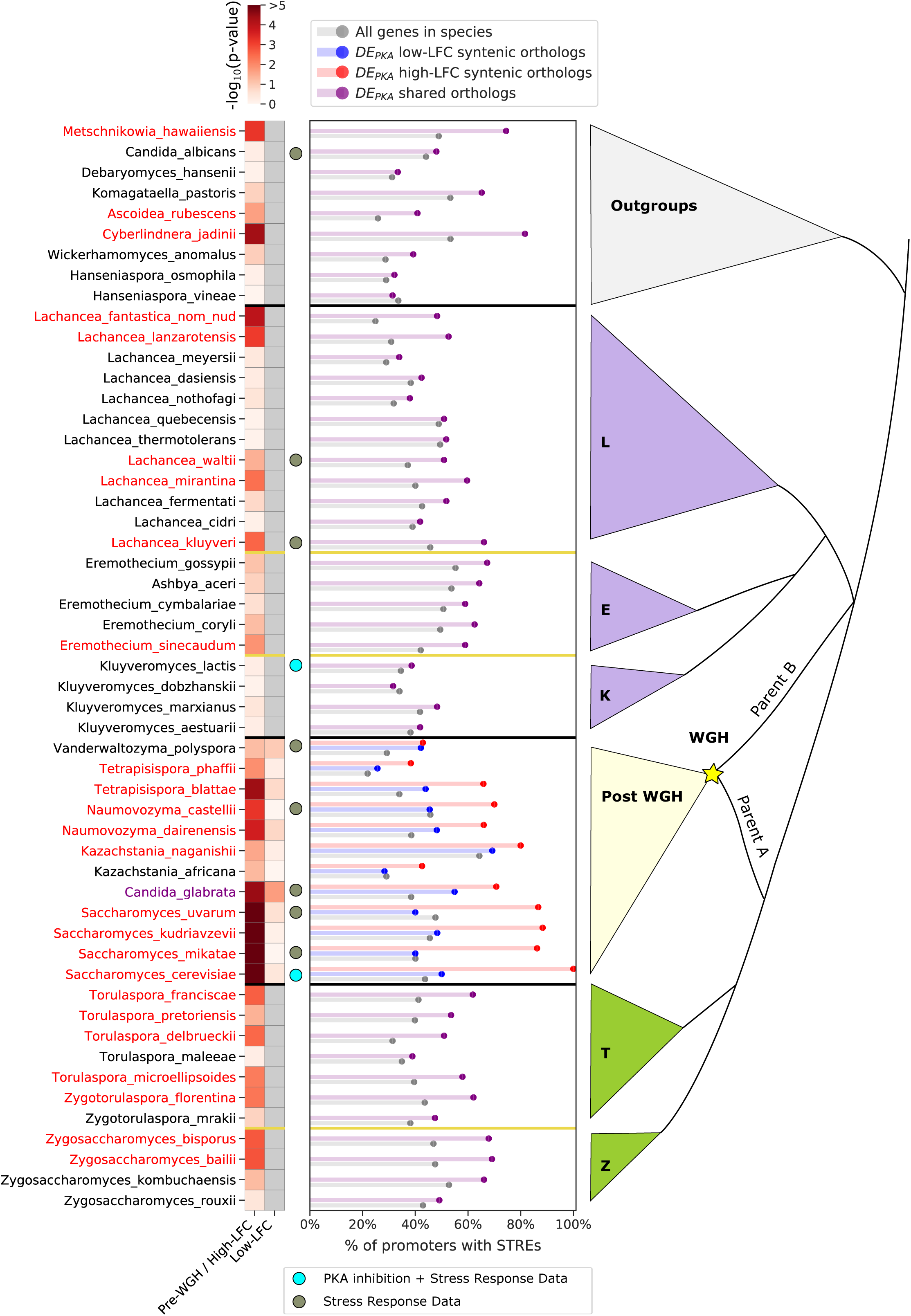
There is enrichment for STREs in the promoters of the syntenic orthologs of the DE_PKA_ high-LFC orthologs, as well as in the ZT branch, indicating that STREs arose in the promoters of DE_PKA_ high-LFC orthologs both after and prior to the WGH. The barplot shows the percentage of promoters with 1 or more STREs in all genes in a species (grey) or in a subset consisting of the orthologs of DE_PKA_ genes (purple for pre-WGH species and blue and red for syntenic orthologs of the low and high-LFC ohnologs respectively). For this analysis we only include ohnolog pairs which had at least one ortholog in the pre-WGH species analyzed, had no more than 8 species with duplicates in pre-WGH species, and had at least one STRE in the high-LFC ohnolog in *S. cerevisiae* (60 ohnolog pairs). The heatmap shows the p-value (Fisher’s exact test) of the hypothesis that the percentage of promoters with at least one STRE in a given set is different from that percentage for all genes in that species. For pre-WGH species, species names are colored red when p<0.05 for the DE_PKA_ orthologs. For post-WGH species, species names are colored red when p<0.05 for the syntenic orthologs of the DE_PKA_ high-LFC ohnologs. *C. glabrata* has a purple label indicating that p<0.05 for both the DE_PKA_ high and low-LFC ohnologs. Grey dots next to a species indicate that gene expression data is available for stress conditions, and light blue dots are beside *S. cerevisiae* and *K. lactis* which have data for PKA inhibition as well as for stress conditions.

Further analysis of the promoters of the pre-WGH orthologs of this subset of DE_PKA_ ohnolog pairs revealed more enrichment for STREs in the ZT branch (7 out of 11 species) than in the KLE branch (only 6 out of 21 species) (Fig 5 heatmap). In the outgroups, there was a mixed picture, with 3 out of 9 species showing enrichment for STREs. Despite the fact that these data reflected the average of 60 separate sets of orthologs with their own idiosyncratic evolutionary histories, the evidence hints that some of the differential expression we see in DE_PKA_ ohnolog pairs may have arisen in the promoters of pre-WGH orthologs in the ZT branch prior to the WGH.

Taken together this evidence paints a picture in which the STRE, which drives gene expression in response to PKA inhibition via Msn2, seems to have arisen either before or after the WGH in the promoters of the ancestors of differentially expressed ohnologs (Fig 6A). We next explored potential specific examples of both scenarios.

**Figure 6:**
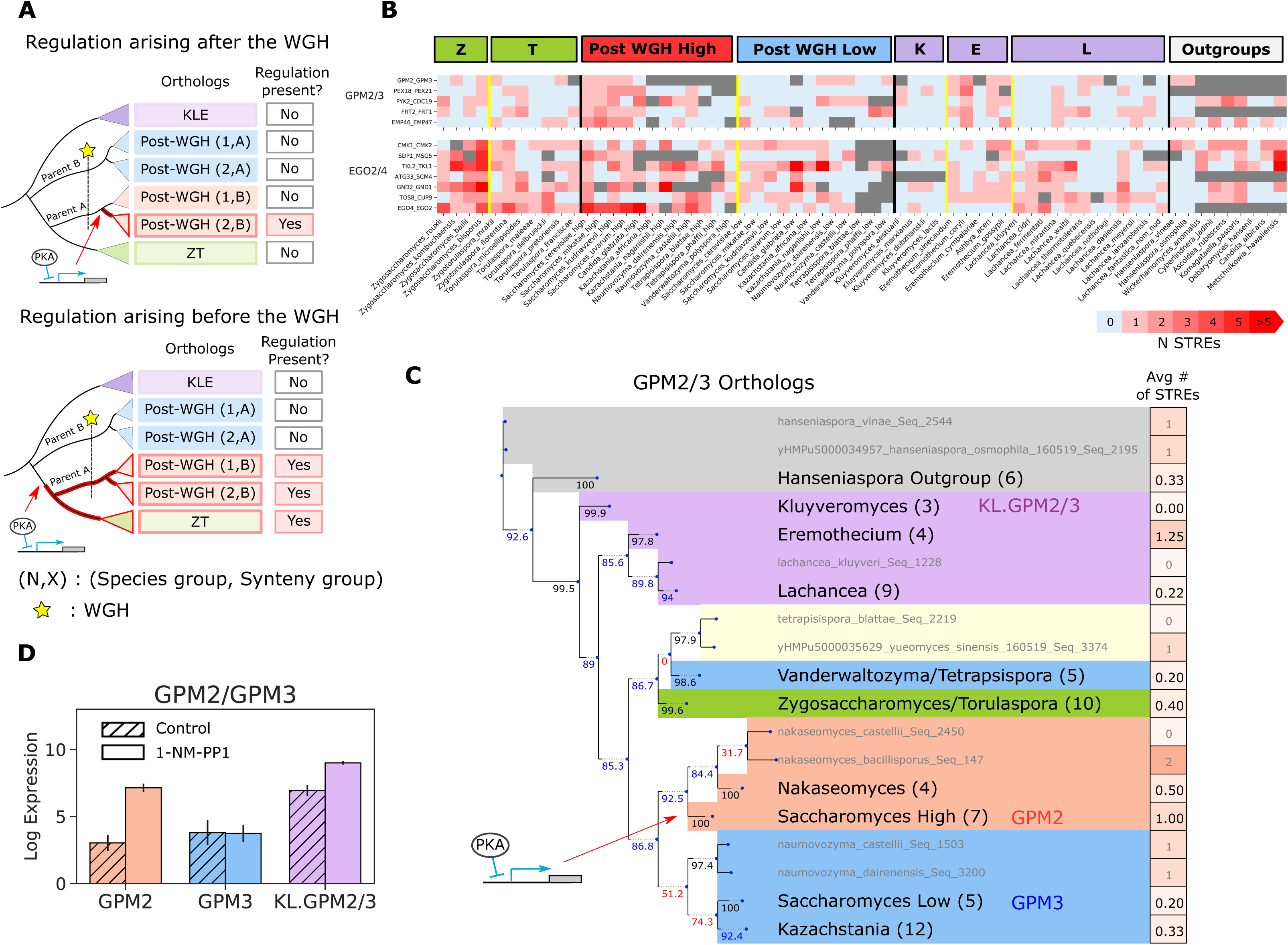
GPM2/3 are an example of a differentially induced pair of ohnologs in which the STRE arose in the promoter of the DE_PKA_ high-LFC ohnolog following the WGH. (A) Schematic depicting scenarios for the appearance of induction in response to PKA inhibition both after (top) and before (bottom) the WGH. (B) Number of STREs in the promoter for two clusters of ohnolog pairs in various species. Grey indicates that no ortholog was found. Data from pre-WGH species and outgroups were used as input to the hierarchical clustering algorithm (see Materials and Methods). (C) Phylogeny of GPM2/3 orthologs alongside average number of STREs in the promoter. Grey text indicates a single ortholog, and black text indicates a group of orthologs with the number of orthologs in that group in parentheses. The putative point at which the STRE was gained is indicated by the red arrow. Bootstrap support values (see Materials and Methods) for each branch point are written to the left of the branch point below the branch and colored black if the support value was above 95, blue if it was between 80 and 95, and red if it was below 80. Promoters are defined as 700 base pairs upstream of the start codon. Shading represents different groups of species; blue = post-WGH, syntenic ortholog to low-LFC ohnolog; red = post-WGH, syntenic orthologs to high-LFC ohnolog; yellow = post-WGH, synteny not determined; green = pre-WGH ZT species; purple = pre-WGH KLE species; grey = outgroups. (D) Average rlog expression with and without 3µM 1-NM-PP1 in GPM2, GPM3, and in their shared ortholog in *K. lactis*, KL.GPM2/3.

### GPM2 and GPM3 illustrate a case in which STREs might have been added following the WGH in the ***Saccharomyces* genus.**

To identify specific promoters with a clear evolutionary signal, we clustered the subset of 60 DE_PKA_ ohnolog pairs described above based on the numbers of STREs in the promoters of their orthologs across 32 ZT and KLE branch species and 9 outgroups (Fig S15). As expected, there is much variation given that the STRE is only a partial surrogate of PKA dependence. However, a few clusters highlight possible examples of WGH paralogs acquiring differential induction under PKA inhibition via distinct evolutionary trajectories. Specifically, one cluster of 5 genes was characterized by low numbers of STREs in the promoters of orthologs from both the ZT branch and the KLE branch (Fig 6B top). This pattern is expected if the STRE and ensuing differential expression would have arisen in the ancestor of the high-LFC ohnolog following the WGH (Fig 6A top).

GPM2/3 provides an example of this situation (Fig 6C, S16). GPM2 and GPM3 were identified in yeast as homologs to the phosphoglycerate mutase enzyme, GPM1, which is part of the glycolysis and gluconeogenesis pathways. GPM2 has been shown to be important for growth under the respiratory carbon sources glycerol and sorbitol, and seems to play a role in the maintenance of cellular membrane stability (Gsell et al., 2013). A specific function for GPM3 has not yet been identified. In our dataset, GPM2 mRNA levels increase substantially under PKA inhibition (5.16 ±0.52 LFC), while GPM3 maintains a stable (albeit low) expression level (3.79±0.87 rlog counts, -0.26±0.75 LFC) (Fig 6D).

To explore the evolutionary history of STRE binding sites in GPM2 and GPM3, we used protein sequence similarity to construct a phylogenetic tree for orthologous yeast proteins and traced binding sites in promoters associated with their coding sequences. We also used syntenic arrangement of the genes surrounding each paralog to identify whether the genomic context for each of the post-WGH orthologs was more similar to GPM2 or GPM3 (red or blue background respectively in Figs 6C, S16, assigned as per Table S6). While most pre-WGH species contained a shared ortholog for GPM2/3, following the WGH duplicate orthologs (ohnologs) were only retained in the *Saccharomyces* genus. GPM3 was more closely related syntenically and in terms of protein similarity with orthologs from the *Kazachstania* and *Naumovozyma* genus, while GPM2 was more closely related to orthologs from the *Nakaseomyces* genus (including the ortholog of *Candida glabrata*).

In every syntenic ortholog of GPM2 analyzed from the *Saccharomyces* genus there is a single STRE between 480 and 510 bases upstream of the gene’s start codon, and multiple TATA box motifs within 150 bases of the start codon (Fig S16). While there are STREs in 3/6 of the closely related *Nakaseomyces* genus orthologs, their locations are not conserved. Thus, a single STRE binding site appears to have arisen in the promoter sequence of the common ancestor of the GPM2 orthologs from the *Saccharomyces* genus. In addition, this STRE binding site was conserved in the promoters of each of those modern GPM2 orthologs (Figs 6C, S16). This motif was presumably part of what gave rise to differential expression between the two paralogs in that clade.

### EGO2/4 provides an example where gain of STREs in the ZT branch may have given rise to PKA dependence that was conserved in syntenic orthologs following the WGH

Another cluster of 7 ohnolog pairs is characterized by a high number of STREs in the ZT branch and relatively fewer STREs in the KLE branch (Fig 6B bottom). This pattern would be expected if the motif arose in the ZT branch after the two branches split but before the WGH (Fig 6A, bottom). EGO2 and EGO4 provide a clear illustrative example of this evolutionary trajectory (Figs 7A, S17).

EGO2 and EGO4 in *S. cerevisiae* are short proteins (75 and 98 residues respectively) that have no clear orthologs outside of the *Saccharomycetacea*, a group which includes the KLE, ZT and post-WGH yeast species. EGO2 was recently shown to be part of the EGO complex which is important for recruiting Gtr GTPases to the vacuolar membrane for TORC1 signaling (Powis et al., 2015). The biological function of EGO4 and its possible role in TORC1 signaling remain unclear. EGO2 is expressed at intermediate levels (6.09±0.12 rlog counts) during exponential growth and does not appreciably change under PKA inhibition (0.41±0.37 LFC) (Fig 7B left). EGO4 has low expression during exponential growth (4.12±0.04 rlog) and is highly induced (7.16±0.38 LFC) under PKA inhibition. EGO4 is also an Msn2/4 target because its activation following PKA inhibition in an Msn2/4 deletion strain decreases substantially (to 1.45±0.60 LFC) (Fig S18). We could not identify the shared ortholog of EGO2 and EGO4 in *K. lactis*

**Figure 7:**
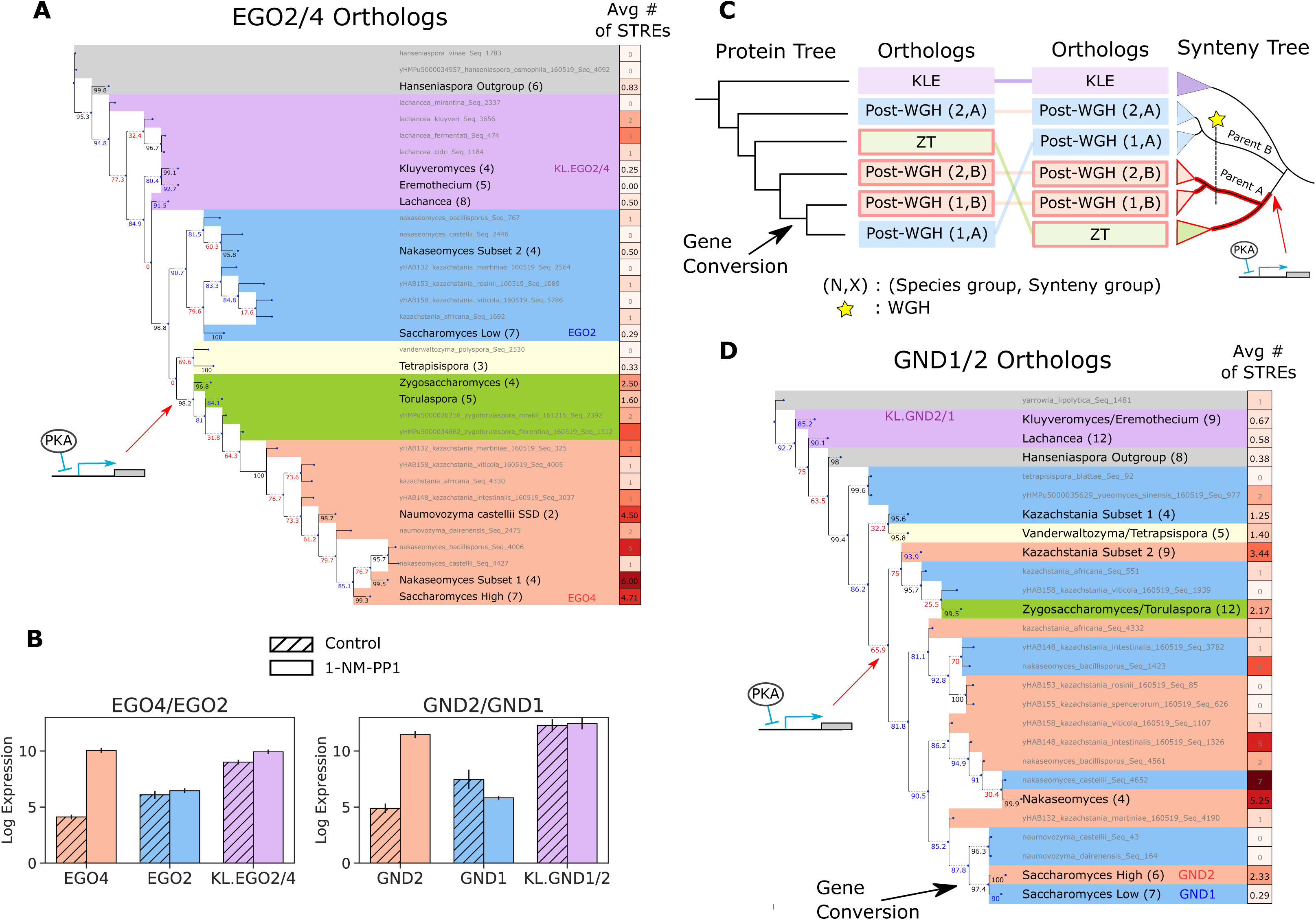
EGO2/4 and GND1/2 are examples of differentially induced ohnolog pairs in which the STRE may have arisen in the ZT branch prior to the WGH. (A) Phylogeny of EGO2/4 orthologs alongside average number of STREs in the promoter. All conventions and nomenclature are as in Fig 6. (B) Average rlog expression with and without 3µM 1-NM-PP1 in EGO2/4, GND1/2 and their shared orthologs in *K. lactis*. (C) Illustration of a protein phylogenetic tree (left) in which a gene conversion has taken place. The synteny tree (right) illustrates the origin of the chromosomal region immediately surrounding the gene conversion. Depending on the size of the region included in the gene conversion, regulatory DNA surrounding the protein following a gene conversion could come from the surrounding region, or from the protein’s original locus. Groups of post-WGH species in the same synteny groups are colored similarly (either blue or red). The WGH is marked with a yellow star. Red outlines indicate conserved regulation by PKA inhibition (e.g. via an STRE in the promoter). (D) As in (A) for GND1/2 orthologs.

We constructed a phylogenetic tree for EGO2 and EGO4, as we did for GPM2/3, using protein sequence similarity to define the structure of the tree and indicated syntenic relationships for post-WGH orthologs with shading (Figs 7A, S17, assigned as per Table S6). The structure of the tree suggested that the protein sequence likely diverged after the ZT and KLE branch split and prior to the WGH, because one branch of post-WGH ohnologs (the one containing EGO4) was more closely related to orthologs in the ZT branch than it was to the other branch of post-WGH ohnologs (the one containing EGO2) (Fig 7A). For this particular set of orthologs the syntenic relationships of post-WGH orthologs corresponded to their phylogenetic relationships based on protein sequence similarity. Such correspondence is not necessarily the case as we will show in a later example.

There were more STREs on average in the promoters of ZT branch orthologs of EGO2/4 (Fig 7A), and all promoters for ZT branch orthologs had at least 1 STRE within 700 bases of the transcription start site, and a TATA box within 300bp of the transcription start site (Fig S17). The location of these motifs was in some cases conserved across species. This pattern was preserved in the promoters of the post-WGH syntenic orthologs of EGO4 and was not generally present in the promoters of the post-WGH syntenic orthologs of EGO2. In the promoters of the KLE orthologs, there were fewer STRE and TATA box motifs and those that were present were scattered more sporadically (Figs 7A and S17). The phylogenetic distribution of this pattern paints a picture in which STREs arose in the promoter of the ancestral EGO2/4 ortholog in the ZT branch before the WGH parental strain (parent A in Fig 1A) separated and were then conserved. The promoter of EGO4, which descended from this lineage, contains some of these conserved STREs. This presumably rendered EGO4 inducible by PKA inhibition (Fig 6A bottom). In this scenario, the promoter for EGO2 would have descended from the WGH parental strain more closely related to the KLE branch (parent B in Fig 1A), which would have had no STREs and would not have been induced by PKA inhibition.

### The STRE in the promoter for GND2 may have arisen in the ZT branch (similar to EGO2/4) but gene conversion obscures its evolutionary history

GND1/2 are an ohnolog pair in the same cluster as EGO2/4 (Fig 6B bottom). These ohnologs represent another example in which the STRE seems to have arisen in the promoter of the common ancestor of the ZT branch and the WGH parental strain more closely related to the ZT branch (Fig 6A bottom). In this case, however, the history of the protein sequence and the gene locus, as determined by synteny, do not coincide. This makes interpretation following the WGH more difficult than in the case of EGO2/4 (Fig 7C).

GND1/2 are two of the 12 enzymes in the Pentose Phosphate pathway (Stincone et al., 2015). Interestingly, our DE_PKA_ set of differentially expressed ohnologs contains three additional ohnolog pairs that are part of the Pentose Phosphate Pathway. In our data, Gnd1 is expressed at intermediate levels during exponential growth and decreases under PKA inhibition (-2.40±0.52 LFC) (Fig 7B right). Gnd2 has low basal expression which increases sharply (7.77±0.40 LFC) following PKA inhibition. The shared ortholog in *K. Lactis* has high expression which doesn’t change under PKA inhibition (0.17±38 LFC). After constructing a phylogenetic tree for GND1/2 and its orthologs using protein sequence similarity we analyzed the promoters of the orthologs. Most (10/12) of the promoters of the ZT branch orthologs of GND1/2 contain at least one STRE in the 700bp upstream of the start codon and a TATA box within 300bp, as was the case with EGO2/4 (Fig 7D, S19). Only 8/21 of the promoters of the KLE branch orthologs have these features, suggesting that the STRE may have arisen after the separation of the ZT branch from the KLE branch.

Determining whether this STRE arose prior to the separation of the WGH parental strain more closely related to the ZT branch (i.e. Fig 6A bottom) requires tracking the evolutionary history of the promoter in post-WGH species. In the case of GND1/2 this task becomes difficult using the protein-based phylogenetic tree because synteny relationships, which correspond to the genomic context of the gene, diverge from relationships based on protein similarity. This divergence is likely due to gene conversion. Gene conversions, in which homologous portions of the genome overwrite one another, occur frequently after interspecies hybridization events, including the WGH in budding yeast (Louis et al., 2012; Marcet-Houben and Gabaldón, 2015). Following a gene conversion in which the syntenic context of the ohnologs does not change, but the protein sequence itself is overwritten by its ohnolog, we would expect the protein-based phylogenetic tree to place the ohnologs close together despite the more distant relationship of the surrounding genomic regions (Fig 7C). Indeed, we see a few cases in the tree of GND1/2 orthologs in which close relationships assigned based on protein sequence similarity do not imply similarity of genomic context in post-WGH genes (indicated by red and blue shading in Fig 7D).

This appears to be the case for GND1 and GND2 in *S. cerevisiae*. These genes and their *Saccharomyces* genus orthologs are more closely related to one another than either are to orthologs from other Pre-WGH species outside of the *Saccharomyces* genus. If we assume that the genomic context of GND1 descends from the WGH parental strain more closely related to the KLE branch (parent B from Fig 1A), then the structure of the protein based phylogenetic tree and the pattern of syntenic similarity implies that the protein sequence of the ancestor of GND1 was overwritten by the ancestor of GND2 without drastically altering the genomic context in a gene conversion event in the ancestor of the *Saccharomyces* genus. There are two conserved STREs upstream of a conserved TATA box in the promoters of the *Saccharomyces* genus orthologs more closely related to GND2 and which are not present in the promoters of the orthologs closer to GND1. While it is possible that the promoter of the ancestral GND2 containing conserved STREs was part of the gene conversion event, and that these binding sites were subsequently lost in the promoter of the converted GND1 ancestor, another more parsimonious scenario is also possible. The regulatory region of the ancestor of GND1, lacking STREs and presumably responsiveness to PKA, could have remained intact during the gene conversion while the protein sequence itself was overwritten by the ancestor of GND1. In this way, the phylogeny of the regulatory region may be separate from that of the protein sequence.

## Discussion

Gene duplication is a major driver of innovation in evolution, and our goal in this study was to better understand the mechanisms that can drive this evolutionary innovation in the special case of an allopolyploidization or Whole Genome Hybridization (WGH). One of the most well-studied examples of a WGH occurred in budding yeast in the branch leading to the canonical single-celled eukaryote, *S. cerevisiae*. This WGH has been implicated as a key factor that enabled the change in metabolic lifestyle from respiratory (or Crabtree negative) to respiro-fermentative (Crabtree positive) that occurred at the same time as the WGH in that lineage (Merico et al., 2007). It is known that the paralogs generated from the yeast WGH (ohnologs) are often differentially expressed under stress conditions. However, the pleiotropic nature of environmental stress makes it difficult to parse the contribution of different interacting stress response pathways to differential expression. By narrowing our study to the PKA pathway, a conserved master regulator of the Environmental Stress Response (ESR), and capitalizing on ways to perturb the pathway specifically, we could meaningfully explore how the signal from PKA was delivered to ohnologs that are differentially expressed. Additionally, in comparing the transcriptional response to PKA inhibition between the pre-WGH species *K. lactis* and the post-WGH species *S. cerevisiae*, direct control of PKA ensured that changes we observed were the result of changes downstream of this particular pathway rather than changes in the stress sensing machinery and the crosstalk between pathways.

Our data established a strong enrichment for ohnologs from the yeast WGH in the set of genes activated by PKA inhibition in *S. cerevisiae* and revealed that many of these ohnologs had differential expression, with one member of the pair showing high induction by PKA inhibition and the other remaining insensitive to PKA. This set amounts to a large proportion (almost 1/6^th^) of the retained ohnologs in *S. cerevisiae*. Our investigation of the transcriptome of *K. lactis* in response to PKA inhibition, and our further analysis of publicly available gene expression data from a number of budding yeast species in response to stress conditions that approximated PKA inhibition revealed that the orthologs of these differentially expressed ohnologs in various WGH-species had low induction and high basal expression. These data suggested that insensitivity to PKA is likely to be the ancestral state for many of these differentially expressed ohnologs (Roy et al., 2013; Thompson et al., 2013; Tsankov et al., 2010). To explore the evolutionary timeline for the emergence of PKA dependence for the ohnologs that were induced by PKA inhibition, we looked for bioinformatic signals in the promoters of differentially expressed ohnologs and their orthologs across over 70 sequenced budding yeast genomes. We found compelling evidence that the STRE binding site, which is strongly linked to activation by Msn2/4 under PKA inhibition, was gained in the ZT branch prior to the WGH in some cases and arose following the WGH in others. The examples of gene promoters we further analyzed as examples of these scenarios highlighted additional interesting evolutionary issues.

Given the immense challenge faced by natural selection to maintain redundancy despite mutations, understanding how duplicated genes persist is one of the important questions in understanding evolution. Many hypotheses have been put forward, including the “balance hypothesis” (Papp et al., 2003), which posits that paralogous genes that are dosage dependent (i.e. members of essential complexes) are likely to be retained following a WGD or WGH, since the loss of one copy would cause a substantial fitness defect. This is likely to be the case for the large contingent of ribosomal ohnolog pairs that are simultaneously repressed under PKA inhibition in our data. Other evolutionary forces that work to generate functional divergence in paralogs include escape from adaptive conflict, sub-functionalization, and neofunctionalization (Des Marais and Rausher, 2008; Hittinger and Carroll, 2007). In an autopolyploidization event, or WGD, however, one still must explain how such mechanisms are deployed. Because in a WGD the two copies of a gene are initially identical, one must invoke either very strong selection for advantageous functional divergence or easily accessible mutations that provide this divergence for it to arise before deleterious mutations have a chance to break one of the two identical paralogs.

An allopolyploidization, or WGH, on the other hand provides a simpler explanation for the origin of functional divergence in paralogous genes. Two paralogous genes might have functionally diverged during the time they were separated in different lineages, each adapting to its own specific environment. We envision this to be the case for EGO2/4 genes, which arrived in *S. cerevisiae* with different regulation (one downstream of PKA and one not) and different protein phylogeny. GPM2/3 may also have experienced functional divergence prior to the WGH, but it appears that differential regulation in response to PKA (in the form of STREs in the promoter) did not arise until afterwards. However, this simple story is often greatly complicated by the occurrence of gene conversion following WGH events when homologous genes overwrite all or part of their paralogs either soon after the polyploidization event or in the following millennia (Louis et al., 2012). It has been proposed that the functionally divergent ohnolog pair GDH1 and GDH3 experienced a gene conversion following the WGH(Campero-Basaldua et al., 2017). Our investigations of GND1/2 indicate that they might constitute another example of gene conversion.

Understanding the implications of gene conversion following a hybridization (versus a duplication) event is therefore crucial. In a hybridization event, while the protein sequence is homogenized through gene conversion, divergent regulation is likely to remain in the form of any different cis-regulatory elements in the promoters of the paralog pairs that were unaffected by the gene conversion. A corollary of that is the idea that a WGH can provide an environment in which functional divergence could evolve as different ohnologs gain mutations that either enhance their function in conditions under which they are expressed or degrade functions required for conditions under which they are not. This may have been the case for GND1/2. The same reasoning applies if a cis-regulatory sequence is lost or appears de-novo in the promoter of one of two duplicated ohnologs, causing differential expression and paving the way for further functional specialization.

Stronger support for these insights would be provided by measurements of gene expression under direct inhibition of PKA in a range of budding yeast species, including important members of the ZT branch. Such measurements would pave the way for inferring the levels of divergent expression that were present in the ancestral hybrid, and for determining how much arose de-novo in the various post WGH lineages. Another fruitful direction in exploring the evolutionary trajectories that we propose would be to perturb cells further down the PKA pathway at the level of the transcription factor Msn2 in order to avoid possible crosstalk that might exist downstream of PKA. This is technically challenging for Msn2 because its activity is predominantly controlled by its nuclear localization. However technology to optogenetically induce nuclear localization for transcription factors such as Msn2 has recently been demonstrated and might be possible to port into other budding yeast species (Chen et al., 2019).

The STRE motif is a bioinformatic signal with clear weaknesses, including a relatively low information content (just 5 bases) and in imperfect correlation between the signal (the STRE) and the biological phenotype it was meant to detect (induction in response to PKA inhibition). Despite this, we were still able to leverage the high density of sequenced species in budding yeast to draw evolutionary conclusions (Shen et al., 2018), illustrating the usefulness of the budding yeast as a model phylum for studying evolution (Botstein and Fink, 2011). The rest of the known budding yeast will soon be sequenced completing the goals of the y1000plus project (“Y1000Plus,” n.d.) and, unless more extant and ancient budding yeast are later discovered and sequenced, that data will represent all the evidence that remains of hundreds of millions of years of regulatory evolution for those species. Our ability to extract insights into regulatory evolution from that data awaits a better understanding of a few key links in the chain connecting the pathways that sense environmental changes (such as PKA) to gene expression. Breakthroughs at a few levels might soon yield the technology necessary to read bioinformatic regulatory signals in these yeast at a much greater level of precision. A screen of pathway perturbations linked to high throughput gene expression readout, such as single cell or single colony RNA-seq (Jackson et al., 2019; Liu et al., 2019), combined with machine learning approaches would make great strides towards identifying such a bioinformatic regulatory signal for the PKA pathway. This would be strengthened by a tighter link between pathway mutants and transcription factor binding via traditional assays such as ChIP-seq and complementary methods such as the calling card method (Mayhew and Mitra, 2016). It would also be strengthened by incorporating the fuller understanding of the links between transcription factors and the DNA sequences they bind, both in vitro and in vivo (Le et al., 2018; Samee et al., 2019; Zhu et al., 2018). A combination of all such methods will allow us to more fully understand the role that changes in gene regulation play in evolution.

## Supporting information

Supplemental Table 3: Ohnolog pair data

Supplemental Tables 4 and 5: Strains and Plasmids

Supplemental Table 6: Syntenic ohnolog assignment

## Acknowledgements

We thank Nan Hao for providing PKA-AS strains in *S.cerevisiae* and Alexander Johnson and his lab, especially Trevor Sorrells and Chiraj Dalal, for providing helpful advice, feedback and *K. lactis* strains and plasmids. We also thank Patricia Babbitt for helpful discussions and feedback. We thank Snigdha Poddar and Jamie Cate for assistance with the CRISPR gene editing protocol in yeast, as well as Ivan Liachko and Maitreya Dunham for providing the Pan-ARS plasmids used as a backbone for our gene editing in *K. lactis.* We thank Eric Wong, Derrick Lee, Derek Britain, Melanie Sylvis, Gabe Reder and Kyle Fowler for discussions and ideas. Thanks also to Andrew Prokop for assistance in adapting the CRISPR gene editing protocol to yeast and to the entire El-Samad lab for helpful discussions and feedback along the way, especially to Jacob Stewart-Ornstein and David Pincus.

## Author contributions

Study conceptualization and experiment design, B.M.H. and H.E.-S.; Experiments and bioinformatic analysis B.M.H.; Wrote the paper, B.M.H. and H.E.-S.; Supervision and funding, H.E.-S.

## Materials and Methods

### Plasmid and strain construction

All plasmids and strains used in this study are listed in Tables S4 and S5.

The *S. cerevisiae* PKA-AS base strain was obtained from of Nan Hao and Erin O’Shea (Hao and O’Shea, 2011). In addition to the gatekeeper point mutations (M164G, M147G and M165G for TPK1, TPK2, and TPK3 respectively), the strain contained an NHP6A-IRFP nuclear marker which was not used in this study.

The *K. lactis* PKA-AS strains (yBMH132, yBMH078), containing the M222G and M147G mutations for KL.TPK2 and KL.TPK3 respectively, were constructed using a single plasmid CRISPR strategy based on (Ryan and Cate, 2014). We used Gibson assembly to combine Cas9 and sgRNA expression constructs on a backbone with an autonomously replicating sequence from *K. lactis* that allows plasmid replication in a variety of budding yeast species (Liachko and Dunham, 2014). The targeting sequence for that guide was changed using Gibson assembly combining PCR products containing a new guide sequence with the digested backbone from a previously assembled sgRNA plasmid. Donor constructs were designed to have at least 300 bp of homology upstream and downstream from the point mutation, as well as a heterology block consisting of synonymous mutations in the location of the sgRNA target that prevent re-cutting by the Cas9/sgRNA complex as described in (Horwitz et al., 2015a). The donor cassette was printed by SGI-DNA, inc. and integrated into a PUCGA 1.0 backbone.

Auxotrophy for URA in the *K. lactis* WT strain (yLB13a) was made by counterselecting on 5-FOA and confirmed by sequencing.

The transformation for the CRISPR gene editing mutation for the *K.lactis* PKA-AS strain used for RNA-seq and growth experiments (yBMH132) was performed using a standard Lithium Acetate protocol designed for transformations in *S. cerevisiae* based on (Lee et al., 2015) with the following adjustments. 4ml of cells were used for each transformation. The DNA mix contained 5µg Donor DNA PCR amplified and column purified from the Donor DNA plasmid, and 1µg guide plasmid. Colonies were picked after 3-4 days incubation at 30°C.

The transformation for the CRISPR gene editing mutation for the *K.lactis* PKA-AS strain used for the KL.Msn2 nuclear localization experiment (yBMH078) was performed using a the protocol designed for CRISPR/Cas9 transformations in *S. cerevisiae* from (Ryan and Cate, 2014) with the following adjustments. 7.5ml of cells/transformation at OD600 of 0.8 were used to prepare competent cells. Competent cells were washed twice in LATE buffer (100mM Lithium Acetate, 10mM Tris-HCL ph8.0, 0.1mM EDTA ph8.0) prior to resuspending in equal parts LATE buffer (with no PEG 2000) and 50% glycerol prior to freezing at at -80°C. Cells were washed in 1xTE buffer prior to plating on SD-URA and incubated at 37°C for 12-24 hours (instead of 48 hours) followed by 2-3 days at 30°C.

The CRISPR deletion cassettes for *S. Cerevisiae* Msn2/4 deletions were constructed using the plasmids and golden gate protocol from (“Quick and easy CRISPR engineering in Saccharomyces cerevisiae · Benchling,” n.d.) which incorporates in vivo homologous recombination to complete the Cas9/sgRNA expression plasmid per (Horwitz et al., 2015b). A similar set of plasmids was constructed to replace the backbone of the integration vector with the Pan-ARS backbone for use in *K.lactis* using golden gate cloning. Donor DNA for these constructs was constructed using a golden gate strategy to insert the donor sequence into the YTK095 backbone (Lee et al., 2015). The donor sequence was designed to have 60bp homology for *S. cerevisiae* and 300bp homology for *K. lactis* to delete the SC.Msn2/4 or KL.Msn2. proteins respectively. Unlike a deletion cassette strategy, we did not use selection markers and therefore deleted 250bp of the promoter of each protein targeted for deletion in addition to removing their coding sequences in order to prevent spurious expression from an active endogenous promoter. Donor inserts were built using 3 sets of annealed oligos for *S. cerevisiae* or ordered as GeneBlocks (IDT) for *K.lactis*.

Transformations for the CRISPR gene deletions of Msn2/4 in *S. cerevisiae* (yBMH168, yBMH170) and KL.Msn2 in *K. lactis* (yBMH201) were performed using the same standard Lithium Acetate protocol as for yBMH132. The DNA mix for *S. cerevisiae* contained 20ng of BsmBI digested and column purified Cas9/sgRNA expression vector, 40ng of EcoRV digested and column purified sgRNA insertion vector for each mutation (Msn2 and Msn4), and 400ng PCR amplified and column purified donor DNA for each mutation. The DNA mix for *K.lactis* contained 100ng BsaI digested and column purified Cas9/sgRNA expression vector, 200ng of EcoRV digested and column purified sgRNA insertion vector, and 2µg PCR amplified and column purified donor DNA for each mutation.

Following verification of CRISPR point mutations and deletions by sequencing, the Cas9-sgRNA plasmids / expression vectors were removed using counterselection on 5-FOA.

Plasmids for Msn2(C649S) and KL.Msn2(C623S) fluorescent reporters were constructed using restriction digestion and ligation of PCR products. The point mutations that ablate DNA binding for these transcription factors were made using quick change mutagenesis. In addition to containing an mCherry fluorescent reporter for their endogenous Msn2 transcription factors, each strain carried a Venus fluorescent reporter for the Msn2 transcription factor from the opposite species which was not analyzed for this study.

Transformations for the *S. cerevisiae* Msn2 nuclear localization strains (yEW051 and yEW052) were done using a similar lithium acetate protocol as for yBMH132 except for the following variations. An initial amount of 2ml of cells at OD600 of 0.6 were used, the pellet was washed and resuspended in LATE buffer and 30 µl of cell resuspension was combined with 2µl salmon sperm DNA, 120µl 50% PEG-3350, 30µl LITE and 2-5µg digested integration plasmid in 10µl water. Following heat shock cells were washed with 10mM Tris-HCL ph8.0, 0.1mM EDTA ph8.0 (TE) buffer and plated on selective media.

Transformations for the *K. lactis* Msn2 Nuclear localization markers were performed using an electroporation procedure based on that described in (Kooistra and Steensma, 2003) with the following variations. Initially 50ml of OD 0.8 cells were used, wash and DTT buffer volumes were halved, and the volume of final resuspension in electroporation buffer was 240µl. For each transformation, 60µl resuspended cells, originating from about 12.5ml OD 0.8 cell suspension, were mixed with 5µl ssDNA, and 10-20µl DNA mix prior to electroporation, recovery, and plating. DNA mix consisted of 1.5µg cut and column purified integration plasmids in water.

### Growth Experiments

Yeast were picked from freshly streaked plates and grown overnight in YPD to saturation. In the morning they were diluted to OD 0.2 in deep well plates shaking at 900rpm and 30°C in an INFORS-HT Multitron shaker for 1.5 hours. 100µl cells were then combined with 100µl 2x perturbation media containing 1-NM-PP1 (for a final concentration of 3µM) or DMSO in a 96 well glass bottom culture plate (Corning 3904). OD600 measurements were collected using 10 flashes and a settle time of 100ms on a TECAN SPARK10M plate reader every 20 min using Spark Control V2.2 software. A humidity cassette was used with no lid on the plate. Temperature control was set to 30°C, and the plate was set for continuous double orbital shaking (2.5mm, 108rpm). Growth data was analyzed using custom python scripts located at https://github.com/heineike02/AHN_FlowTools.

### Microscopy

Yeast were grown overnight to saturation in 4ml SDC and diluted to approx. OD 0.05 in the morning to get OD 0.2 after 5 hours (6.5 hours for K. Lactis) assuming a lag time of 90 min for both species and a doubling time of 115 for *S. cerevisiae* and 130 min for *K. lactis*. 96 well glass-bottom plates (Brooks Life Science Systems MGB096-1-2-LG-L) were prepared for imaging by coating with 0.25mg/ml concanavalin A (Sigma-Aldrich C2010, resuspended in distilled water) (ConA) for 30 min and washed twice with SDC prior to adding cells. Cells were sonicated gently (amplitude 1 for 3s using a Misonix S-4000 sonicator with a 1/16in microtip P/N #418) to separate clumps and incubated in appropriate wells for 30min. Following incubation, cells were washed twice with 100µl SDC leaving 100µl of SDC in the well above immobilized yeast cells. After 2-3 initial images 100µl of 2x perturbation media (SDC plus 1-NM-PP1 for a final concentration of 4µM, or a DMSO control) was added to the plate. Cells were then imaged every 2-3 minutes until the end of the experiment.

Widefield microscopy images were taken on a Nikon Ti-E inverted scope, with an incubation enclosure set to 30°C, and mercury arc-lamp illumination using RFP filters and dichroic mirror from the Chroma 89006 ET-ECFP/EYFP/mCherry filter set. The microscope’s perfect focus system was used to maintain focus on the samples throughout the experiment. The microscope was controlled with micro-manager software version 1.4.17 using the High Content Screening Site Generator Plugin (Edelstein et al., 2014). Images were taken with an Andor 512 pixel EMCCD camera (897 iAxone DU-897E) using a 40x objective (Nikon Planfluor 40x/0.75) and 1.5x optical zoom.

Image analysis was conducted using custom MATLAB scripts located at https://github.com/heineike02/image_analysis. Briefly, RFP images are background subtracted, smoothed, and cells are identified using the Lucy-Richardson deblurring algorithm (MATLAB function deconvlucy) with 5 iterations based on an image of a single typical cell of the appropriate species. For each identified cell, nuclear localization is calculated as the mean intensity of the top 5 brightest pixels to the median pixel intensity.

### RNA sequencing

Cells were grown overnight at 30°C in YPD to saturation, and then diluted to obtain a 30ml of cells with an OD600 of 0.5 in 4-6 hours assuming a lag time of 90 min and a doubling time of 90 min for *S. cerevisiae* and 110min for *K. lactis*. The culture was then divided into two 12ml treatment and control cultures and 3.6µl of 10mM 1-NM-PP1 (for a final concentration of 3µM) or DMSO was added respectively. Each culture was incubated while shaking at 250RPM and 30°C and split into two 5ml aliquots prior to spinning down at 3850RPM (331rcf) an Eppendorf 5810R benchtop centerfuge for 3min. After spinning down, supernatants were poured out and remaining supernatant was aspirated off with a P1000 pipette. Cells were then flash frozen in liquid nitrogen and stored at -80°C. RNA extraction was performed using the hot acid-phenol extraction protocol of (Solís et al., 2016) with the following changes. Initial cell volume was 5ml instead of 1.5ml and thus the initial spin for collecting the cells was done for 3min at 3850RPM (331rcf) instead of 30s at 13000RPM (15871rcf). Acid-phenol:chloroform:isoamyl alcohol (IAA) (125:24:1), pH 4.5 (AM9722) was used instead of pure Acid-Phenol because the chloroform aids in the separation of nucleic acid from proteins and lipids, and the IAA prevents foaming. The spin following heat incubation was performed at room temperature instead of at 4°C. A second 400µl chloroform wash was included prior to removing the aqueous phase from the phase lock tubes. 22µl of 3M NaOAc, pH5.2 was added instead of 30µl to precipitate the DNA. Ethanol Precipitation was done overnight instead of for 30 min.

Total RNA was aliquoted and stored at -80°C. Prior to library preparation, total RNA concentration was measured estimated using a Nanodrop 2000c spectrophotometer and screened for quality by electrophoresis. For electrophoresis, a formamide based loading dye was used to run samples on a 1.2% agarose gel in TBE buffer. Select samples were also checked for quality using an Agilent 2100 Bioanalyzer with an RNA 6000 Pico chip. 3’ Sequencing Libraries were prepared using the Lexogen QuantSeq 3’mRNA-Seq Library Prep kit FWD using dual indices for each sample. Select libraries were checked for quality on the Bioanalyzer using a High Sensitivity DNA chip. Library concentrations were calculated using a Qbit 2.0 Fluorometer (Invitrogen), and 2.65 ng per sample were pooled and sequenced on an Illumina Hiseq 4000 sequencer to an average depth of between 245,000 and 5.6 million reads per sample (median 2.53 million). At least 3 replicates were collected for each sample and condition.

Sequencing data was trimmed, aligned, and checked for quality using the Bluebee genomics analysis pipeline. *S. cerevisiae* samples were run using the “FWD S. cerevisiae (R64) Lexogen QuantSeq 2.2.3” protocol, and *K. lactis* samples were run using the “FWD K. lactis (ASM241v1) Lexogen Quantseq2.2.3” protocol. The counts generated with these pipelines used the saccharomyces_cerevisiae_R64-2-2_20170117 GFF for *S. cerevisiae*, and many of the 3’ sequencing reads fell into unannotated 3’UTR regions. We therefore created a modified GFF file which included 3’ UTRs from (Nagalakshmi et al., 2008). We had a similar issue for the *K. lactis* reads, but as there are no definitive studies annotating the *K. lactis* 3’UTR, we created a *K. lactis* GFF with 400bp extensions for each annotated gene serving as estimated 3’UTR regions. We then recalculated gene counts for each species using htseq in intersection-nonempty mode. Custom scripts and the updated GFF files used for these calculations are available at https://github.com/heineike02/UTR_annotation and https://github.com/heineike02/rna_seq_processing.

Gene count data was processed to yield Log Fold Change and pValue estimates using the DESeq2 (Love et al., 2014) package in R (Bioconductor). Log Fold Change and pValue for PKA-AS strains (both WT and ΔMsn2/4 or ΔKL.Msn2) in each species were calculated using all replicates in the respective strain +/-drug at 50 min. Samples which had no counts for any condition were removed. Rlog values for each replicate in *S. cerevisiae* and *K. lactis* were calculated using a DESEQ call that included data from 59 and 35 samples respectively which were sequenced in the same run. Mean rlog data for each condition was calculated by averaging rlog values across replicates. These samples included the WT +/- drug and AS +/- drug experiments as well as experiments with Msn2/4 deletion and Rph1/Gis1 deletion mutants. To calculate distributions of rlog values for all genes in *S. cerevisiae,* 443 orfs classified as dubious in the saccharomyces_cerevisiae_R64-2-2_20170117 GFF were filtered out. Custom scripts for further RNA sequencing analysis and enrichment are located at https://github.com/heineike02/yeast_esr_expression_analysis

### GO term enrichment analysis

The GO Slim database used for analysis was downloaded from SGD on 20180412 (SGD Project, 2018), and only Biological Process terms were analyzed. Fisher’s exact test was used to test the hypothesis that the proportion of genes with a particular GO Slim term in a given set is greater than would be expected given the distribution of that term in a background set. The background set for the genes only activated or repressed in *S. cerevisiae* was all genes in *S. cerevisiae.* The background set for the sets of genes involving activation or repression in *K. lactis* was the set of all genes in *S. cerevisiae* which contain orthologs in *K.lactis*.

### Ohnolog and ortholog assignment

The ohnolog dataset for *S. cerevisiae* was downloaded from the Yeast Genome Order Browser (YGOB) website on 10 October, 2017. Fisher’s exact test was used to test the hypothesis that the proportion of ohnologs in a given set is greater than would be expected given the distribution of ohnologs in the background set. The same test sets and background sets were used as for GO term enrichment analysis.

Ohnolog sets for other species were identified using the “pillars” database downloaded from the Yeast Genome Order Browser on 08 September, 2016.

Orthology assignments between *S. cerevisiae* and *K. lactis* were generated using the YGOB pillars database.

We used orthology assignments from the fungal orthogroups (Wapinski et al., 2007) for comparisons of microarray data from (Roy et al., 2013; Thompson et al., 2013) when present. That database contained no orthology assignments for *V. polymorpha*, so orthology assignments from *S. cerevisiae* to *V. polymorpha* were generated using YGOB pillars.

The gene names used for *K. lactis* and *N. castellii* on the fungal orthogroups website and in those microarrays was inconsistent with the standard gene names which are used by YGOB. We therefore generated a mapping for mismatched gene names between the two databases using protein similarity based on the genome sequences provided by each study. To generate a similarity score between candidate proteins, we used the pairwise2.align.globalms function from biopython with the following parameters: match_points=1, mismatch_points=-1, gap_open=-0.5, gap_extension = -0.1. We then took as candidates the top 5 genes above a threshold (*K. lactis*: 138, *N. castellii:* 115). We kept the top candidate and any lower scoring candidates as long as there was not a drop in 10 points between that candidate and the next highest scoring gene. We generated similar mappings going the opposite direction from the YGOB genename to the fungal orthogroup genename for *N. castellii, K. lactis,* and *S. mikatae* (threshold scores of 120,138, and 130 respectively), and a mapping from the *K. lactis* standard gene name to the gene identification numbers assigned in (Shen et al., 2018) using a threshold score of 110.

To identify syntenic orthologs of DE_PKA_ genes within post-WGH species in order to analyze their promoters, we used the YGOB webtool with a window of +/-8 genes.

To determine orthology relationships for genes from (Shen et al., 2018), we used orthogroups as defined by that paper’s orthomcl.clusters.txt file.

To identify syntenic orthologs within the post-WGH species for our three example ohnolog pairs (EGO2/4, GND1/2, and GPM2/3) we used YGOB to generate a syntenic alignment with a window of +/- 8 genes around each example gene. We extracted the surrounding genes from the sequenced genomes for all species and assigned orthology for the surrounding genes based on presence in the same orthogroup as *S. cerevisiae* and *K. lactis* genes. We then manually curated synteny based on the pattern of orthologs present (Table S6).

### Microarray data processing

Gene expression datasets from microarrays measuring the response to stress conditions in various budding yeast species from (Thompson et al., 2013) (GSE36253) and (Roy et al., 2013) (GSE38478) were downloaded from the NCBI Gene Expression Omnibus (https://www.ncbi.nlm.nih.gov/geo). The data represented Log Fold Change of intensity for the experimental condition divided by the control condition. Where there were multiple datapoints for the same gene name, the median value was used. We took the median value of technical replicates. To compare data across species, we combined all data for all conditions for a given species into a single dataset, and then used the mean and standard deviation to center and normalize all data for that species.

Microarray data estimating raw expression by comparing mRNA from cells collected during exponential growth on rich media to genomic DNA from (Tsankov et al., 2010) (GSE22193) were downloaded from the NCBI Gene Expression Omnibus (https://www.ncbi.nlm.nih.gov/geo). The median of all data that had more than one spot per gene name was used for each gene name. Data for replicates were quantile normalized (Bolstad et al., 2003) using the implementation of (https://github.com/ShawnLYU/Quantile_Normalize), and then the median of the replicates was used for each gene name. Before comparing data across species, the data were mean-centered and normalized by the standard deviation.

### Identification of PKA targets

Genes were classified as activated and repressed by PKA inhibition for GO term and Ohnolog enrichment analysis in both *S. cerevisiae* and *K. lactis* as illustrated in Figure S2. A minimum p-value was defined - log10(pvalue)>ymin where ymin=1.5. An LFC threshold that depended on the p-value was chosen (so that genes with high p-values required a larger LFC in order to be members of the set). The threshold lines are described in (LFC, -log10(pvalue)) coordinates by (x1, y1) and (x2, y2) for both *S. cerevisiae* and *K. lactis*. The coordinates that described the LFC threshold for genes activated by PKA inhibition were (x1_act, y1_act) = (2.0, 15.0), (x2_act, y2_act) = (2.5, 0.0), and those describing the LFC threshold for genes repressed by PKA inhibition were (x1_rep, y1_rep) = (-2.5, 0.0), (x2_rep, y2_rep) = (-2.0, 15.0) and ymin = 1.5.

### Differential expression definition

In *S. cerevisiae* we identified differentially expressed genes by first identifying ohnolog pairs in which one member of the pair was either activated or repressed more than twofold (LFC > 2.0 for or LFC<-2.0, respectively) with a log10(pvalue) less than -1.5. We then retained from this set members where the other ohnolog in the pair did not change in the same direction, specifically having LFC less than 1.5 or LFC greater than -1.5 respectively. We also further required that the difference of estimated LFC between the two paralogs be greater than 2.0 in LFC.

In other species, and for the set of genes differentially expressed under PKA related stress conditions, we identified differentially expressed genes by first defining an estimated Log Fold Change by averaging across the 3-5 (depending on the species) PKA inhibition related conditions (LFC_est_). As in the *S. cerevisiae* RNA-seq data, we first identified ohnolog pairs in which one member of the pair was activated (LFC_est_>1.8) and retained pairs in which the other member was not activated (LFC_est_ < 1.6). We also required that the difference in LFC_est_ between the activated and non-activated ohnolog was greater than 1.8. These thresholds were chosen because they yielded a good overlap between 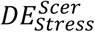 and DEPKA.

### Promoter extraction

Promoter sequences were taken to be the 700bp upstream of the start codon (except when the scaffold contained less than 700bp in which case all bases were used). *S. cerevisiae* promoter sequences were taken from SGD. Promoter sequences for *K. lactis* were extracted gene by gene using NCBI E-utilities. Promoters for YGOB species used in motif analysis for genes that were targets of PKA related stress conditions (Fig 4D), as well as for counting STRE motifs in post-WGH species in figures 5 and 6B were extracted from the genomes for those species on the YGOB website. Promoters for genes in pre-WGH species in figures 5 and 6B and for all species in figures 6C and 7 were extracted from the genomes genomes published in (Shen et al., 2018) using custom scripts available at https://github.com/heineike02/y1000plus_tools,

To analyze promoters of the orthologs of DE_PKA_ genes in 32 pre-WGH *Saccharomycetaceae* species and 9 outgroups, we first identified orthologs using orthogroups calculated in (Shen et al., 2018). We removed any ohnolog pairs that did not contain any STREs in the high-LFC ohnolog reasoning that the high-LFC ohnolog in those pairs were induced through a mechanism that doesn’t require the STRE. We also removed pairs that had that had no orthologs in pre-WGH species (such as USV1/RGM1). Finally, we removed pairs that had more than one ortholog in more than 8 pre WGH species. This was to avoid ohnolog pairs that were present as duplicates before the WGH (such as the hexose transporters) and ohnolog pairs whose pre-WGH orthologs may have undergone a duplication following the WGH, but still ancient enough to be present in several species (such as SNF3/RGT2). This left 60 of 91 DE_PKA_ ohnolog pairs.

### Motif Enrichment

De novo motif identification was conducted on indicated sets of promoters using the DREME algorithm from the MEME suite (Bailey, 2011) with default promoters and a background set of all promoters from either *S. cerevisiae* or *K. lactis* as appropriate.

STRE (CCCCT) and TATA box (TATA(A/T)A(A/T)(A/G)) motifs were identified by looking for exact matches within the promoters, and enrichment was calculated using Fisher’s exact test.

### Clustering

In order to identify evolutionary trends in STRE appearance before the WGH, we clustered DE_PKA_ ohnolog pairs based on the number of STREs in the promoters of their orthologs in 32 pre-WGH *Saccharomycetaceae* species and 9 outgroups. We created a distance matrix for the 60 ohnolog pairs by using the correlation distance between two rows with missing values removed. Rows were then hierarchically clustered (scipy.cluster.hierarchy) using the ‘average/UPGMA’ linkage method. Separate clusters were defined using the fcluster method with the ‘inconsistent’ parameter set to 1.1.

### Phylogenetic trees

Orthologs for GPM2/3, EGO2/4, and GND1/2 were identified based on orthomcl clusters from (Shen et al., 2018). For EGO2/4 there were two separate clusters so protein sequences for the two clusters were combined for the analysis. A multiple sequence alignment was created using MAAFT version 7.407(Katoh and Toh, 2008) using the E-INS-I algorithm (–genafpair) with –maxiterate 1000. These multiple sequence alignments were then trimmed with trimAL v1.4.rev22 (Capella-Gutiérrez et al., 2009) with the -gappyout option. Evolutionary models were identified as LG+F+I+G4 (for GPM2/3), LG+I+G4 (for EGO2/4), and LG+R5 (for GND1/2) using IQTREE model finder (Kalyaanamoorthy et al., 2017). The same model was selected in three independent runs of the algorithm with w-BIC values above 0.96 in each run. Trees were constructed using IQTREE v1.6.12 (Nguyen et al., 2015). Ten independent runs of IQTREE were conducted on the selected model for each trimmed multiple sequence alignment model using -bb 1000 which calls the UFboot2 algorithm (Hoang et al., 2018) as well as -alrt 1000 to find support values. The tree with the best likelihood by BIC was chosen except when BIC values were equal in which case AIC and likelihood values were also used. For EGO2/4, because there were less variant alignment positions compared with the number of model parameters (77 for both E-INS-I and L-INS-I with 161 parameters to determine branch length and tree structure for 81 sequences), we also built trees using a multiple sequence alignment created with the mafft L-INS-I algorithm (--localpair) to see if the results were robust to variations in the multiple sequence alignment. The chosen model did not change depending on the alignment. The ordering of the clades for that tree did not change depending on the alignment, although the BIC was lower for the original E-INS-I alignment. Support values for EGO2/4 were calculated with the -bnni option to expand the search space for bootstrap trees.

## Supplemental Material

### Extended analysis of promoter sequences

To identify bioinformatic signals associated with the promoters of all genes activated by PKA inhibition in *S. cerevisiae,* we used the DREME algorithm (see Materials and Methods) which identifies short ungapped motifs that are enriched in comparison to a background set of promoters in (Bailey, 2011). Comparing the promoters of all genes activated under PKA inhibition in *S. cerevisiae* against the promoters of all *S. cerevisiae* genes, we identified a motif that strongly resembled the Stress Response Element (STRE, CCCCT) (Fig 4A), the binding sequence for Msn2 and Msn4 (Görner et al., 2002; Smith, 1998). Four of the other five motifs enriched in the promoters of genes activated by PKA inhibition were similar to the STRE or the Post Diauxic Shift element (PDS) motif (T(A/T)AGGGAT) which is itself structurally similar to the STRE (Pedruzzi et al., 2000), while others resembled the TATA-box (TATA(A/T)A(A/T)(A/G)) (E-value 1.3e-3) which is known to be enriched in stress responsive promoters (Basehoar et al., 2004).

Next, we asked how the number and locations of STRE and TATA-box motifs correlated with a gene being responsive to PKA inhibition. Promoters of genes activated by PKA had a larger probability of containing one or more STREs relative to all promoters in *S. cerevisiae* (75.2% vs. 43.9%, p-value 1.5e-16). They also had a notable increase in the average number of STREs per promoter (1.32 vs. 0.62) (Fig 4C). Furthermore, the location of the STREs in the promoters of the genes induced by PKA inhibition had a unimodal distribution with 64.7% of STRE sites found between 100 and 400 base pairs, as opposed to an expectation of 42.9% from a uniform distribution and a 46.1% value when the distribution of STRE locations is compiled for all promoters in the genome (Fig S12A).

The promoters of PKA targets were also enriched for the TATA-box (70.1% with 1 or more TATA box v.s. 57.5% in the promoters of all genes, p-value 2.8e-3) (Fig 4A, S13A). We observed a similar clustering of binding sites between 100 and 400bp upstream of the start codon for TATA box motifs as we saw for STRE motifs (70.7% of TATA-box motifs found between the first 100 and 400 base pairs in promoters of PKA targets versus 57.7% for the promoters of all genes, p-value 2.3e-3) (Fig S13C). Finally, TATA box and STRE motifs are more likely to occur together in promoters of genes activated by PKA inhibition than in all genes (43% of promoters vs. 18%), as expected from the increased percentages of both STREs and TATA boxes in genes activated by PKA inhibition (Fig S14A).

For *K. lactis*, the top hit for promoters of genes activated by PKA inhibition was a motif whose Position Specific Scoring Matrix (PSSM) would be satisfied by an STRE but was closer to a PDS (E-value 4.4e-19) (Fig 4C). Furthermore, the bioinformatic signal for the number of STREs and their location was weaker in *K. lactis* than in *S. cerevisiae* (47.7% of promoters with 1 or more STRE in the promoter in PKA activated genes vs. 34.5% in all genes, p=1.6e-4) (Figs 4B and S12B). In *K. lactis*, as in *S. cerevisiae*, the promoters of genes activated by PKA were enriched for TATA boxes (70.8% with 1 or more TATA box vs. 54.1% in all genes, p=3.8e-4) (Fig S13B). Location clustering in the promoters of genes activated by PKA inhibition for STRE motifs was not apparent in *K. lactis* (Fig S12B), but it was for the TATA box (Fig S13D)

## Supplementary Tables

**Table S1: Go term enrichment for PKA targets.** GO term enrichment was calculated from the GO Slim Dataset downloaded from SGD on April 12^th^, 2018 (SGD Project, 2018) and p-values were calculated using Fisher’s exact test against a background of either all genes in *S. cerevisiae* (for the gene sets containing only genes activated or repressed in *S. cerevisiae*) or all genes in *S. cerevisiae* that contain a *K. lactis* ortholog (for gene sets defined by activation or repression in *K. lactis)*.

**Table S2: Enrichment of ohnologs in genes activated or repressed by PKA inhibition in S. cerevisiae.** Enrichment for ohnologs in *S. cerevisiae* gene sets defined by activation or repression following PKA-inhibition by the gene or by its ortholog *in K. lactis*. P-values were calculated using Fisher’s exact test as in Table S1 against a background of either all genes in *S. cerevisiae* (for the gene sets containing only genes activated or repressed in *S. cerevisiae*) or all genes in *S. cerevisiae* that contain a *K. lactis* ortholog (for gene sets defined by activation or repression in *K. lactis)*.

**Table S3: List of ohnologs pairs with RNA seq data following PKA inhibition.** Ohnologs and related data are sorted into “high” (red shading) and “low” (blue shading) columns based on their LFC. Rows are sorted from highest to lowest LFC for the “high” ohnolog. Purple shading indicates data pertaining to the shared *K. lactis* orthologs. <u>Column Descriptions</u>: DE_pka_act : ohnolog pair is differentially expressed in response to PKA activation with one member activated and the other not activated. Ohnolog pairs in this group are members of DE_PKA_, which is defined such that one member of the pair is activated more than two fold (LFC > 2.0) with a log10(pvalue) less than -1.5, while the other ohnolog in the pair has an LFC less than 1.5. The LFC difference between the two ohnologs is greater than 2.0; DE_pka_rep : ohnolog pair is differentially expressed in response to PKA inhibition with one member repressed and the other not repressed (illustrated in Fig 1D right panel with blue shading). This set is defined similarly to DE_PKA_ but with one member repressed more than two fold (LFC < 2.0) and the other not repressed with an LFC greater than -1.5, and an LFC difference between ohnologs of greater than 2.0; AA%id : percent identity between both ohnologs in *S. cerevisiae*; Length Ratio: ratio of shortest/longest number of amino acids between ohnologs in *S. cerevisiae*; LFC, pvalue, rlog : data from RNA seq experiments (see Methods); act_rep_DEpka: whether the gene is activated (act) or repressed (rep) per the definition of DEpka (LFC>2.0 for activation or LFC<-2.0 for repression, and log10(pvalue)<-1.5.; act_rep_FigS2: whether the gene is activated or repressed for the purposes of our GO enrichment and initial ohnolog enrichment analysis (see Figure S2)

**Table S4: List of strains**

**Table S5: List of plasmids**

**Table S6: Syntenic ohnolog assignment for example proteins.** The YGOB webtool was queried with a window of +/-8 genes to obtain syntenic orthologs for YGOB sequences. For post-WGH species from (Shen et al., 2018), surrounding genes were extracted and assigned orthology based on orthogroup assignments from that work. Orthology to *S. cerevisiae* and *K. lactis* genes from YGOB was used to assign a gene to a given column and then syntenic orthologs were manually assigned based on similarity to the syntenic groups for YGOB species assigned by the YGOB webtool.

## Supplementary Figures

**Figure S1:**
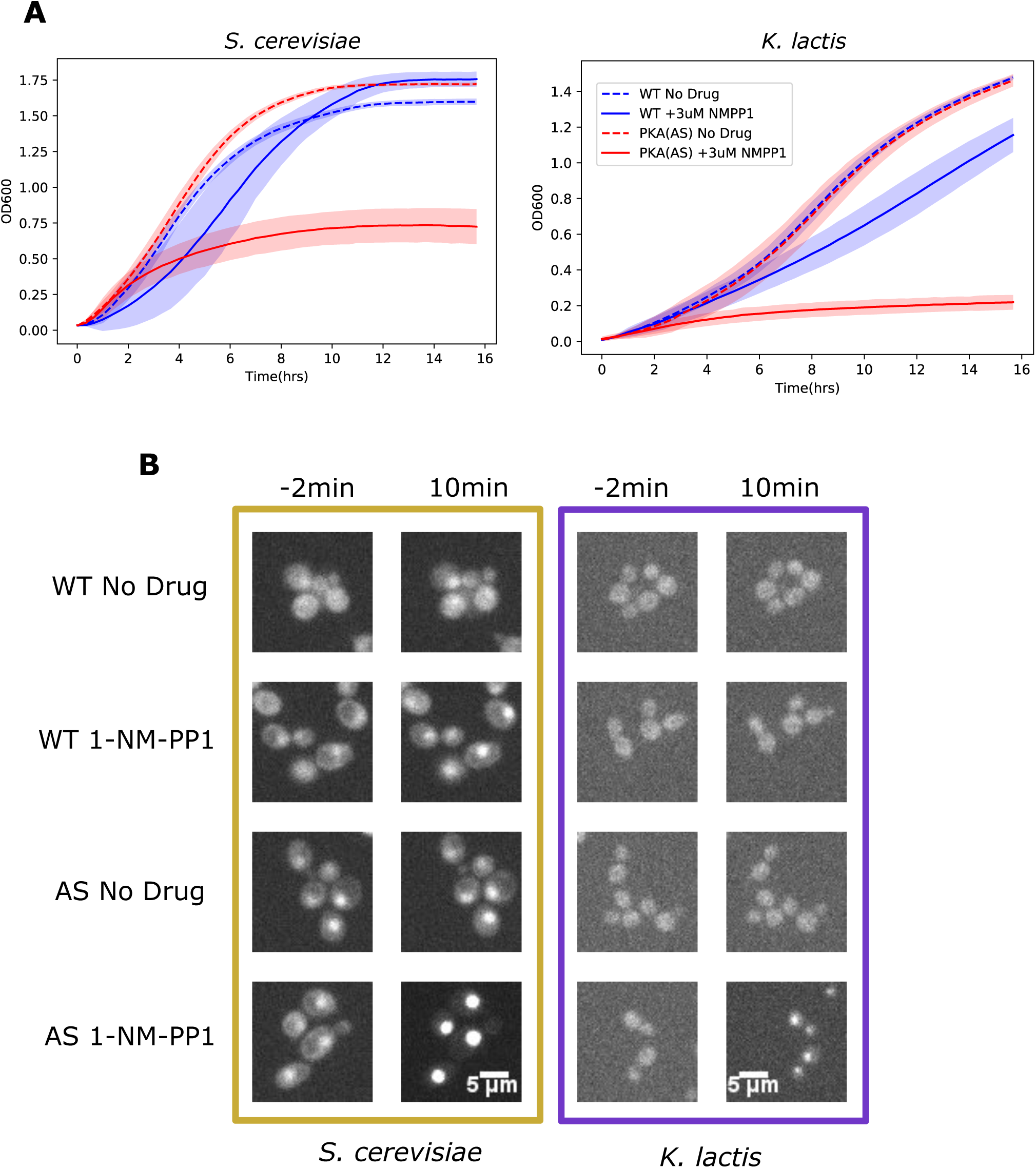
PKA inhibition inhibits growth and causes Msn2 nuclear localization in *S. cerevisiae* and *K. lactis*. (A) WT and PKA-AS *S. cerevisiae* and *K. lactis* strains were grown in YPD in the presence or absence of 3um 1-NM-PP1 and OD600 was measured every 20min. Standard Deviation of at least 4 technical replicates is shown. (B) Selected microscopy images from data in Fig1B. WT and AS strains were grown in SDC and imaged either 2 minutes before or 10 minutes after adding control media or 4uM 1-NM-PP1.

**Figure S2:**
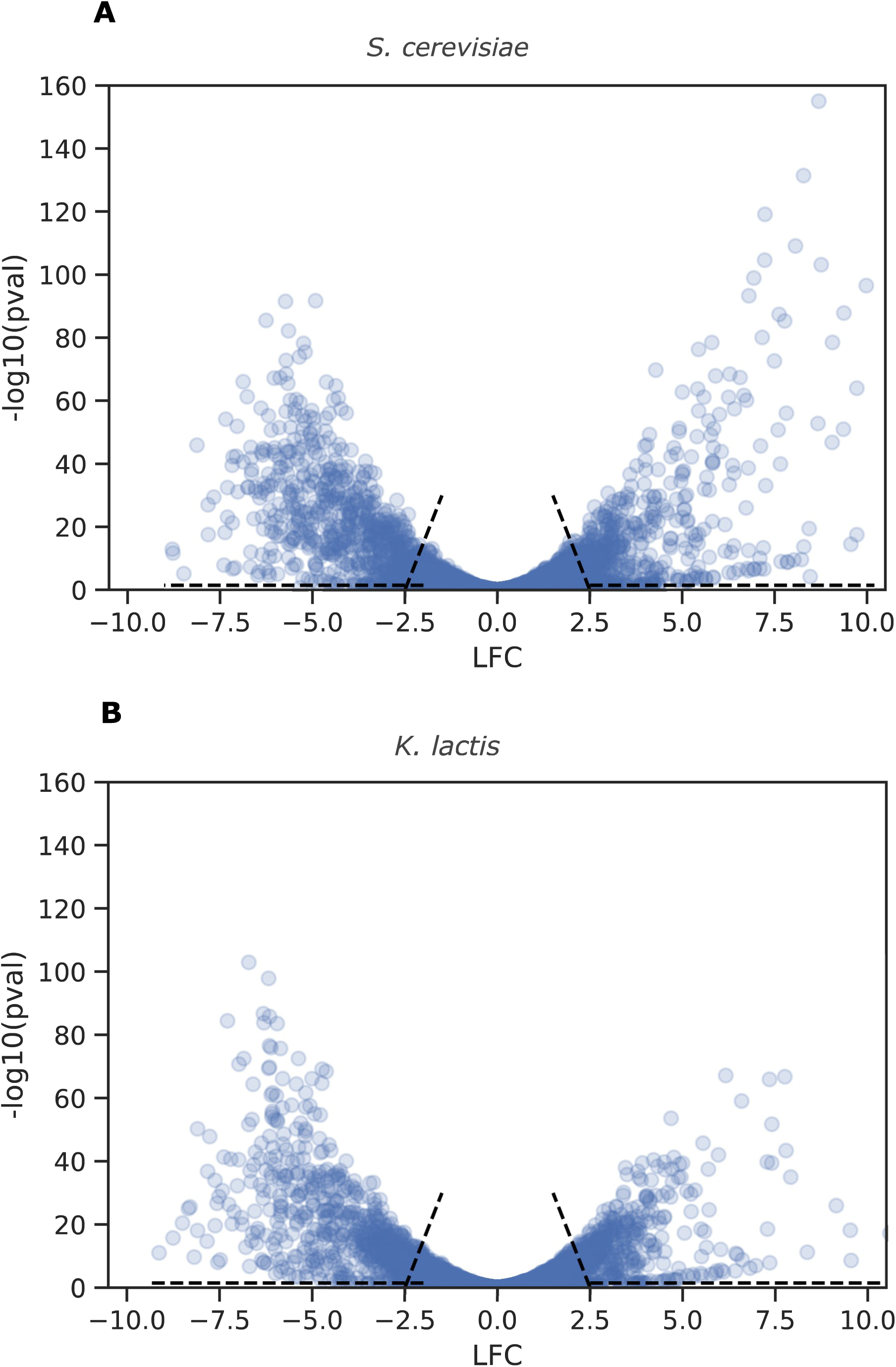
Thresholds defining genes activated and repressed by PKA inhibition in *S. cerevisiae* and *K. lactis*. Volcano plot showing LFC v.s. -log10(p_value) for genes in (A) *S. cerevisiae* and (B) *K. lactis.* Targets of activation or repression in each species were defined as all genes that fell to the right of the lines defined by the points (x1_act, y1_act) = (2.0, 15.0) and (x2_act, y2_act) = (2.5, 0.0), and for which - log10(pvalue) > 1.5. Targets of repression in each species were defined similarly as genes for which - log10(pvalue)>1.5, and which fell to the left of lines defined by the points (x1_rep, y1_rep) = (-2.5, 0.0) and (x2_rep, y2_rep) = (-2.0, 15.0).

**Figure S3:**
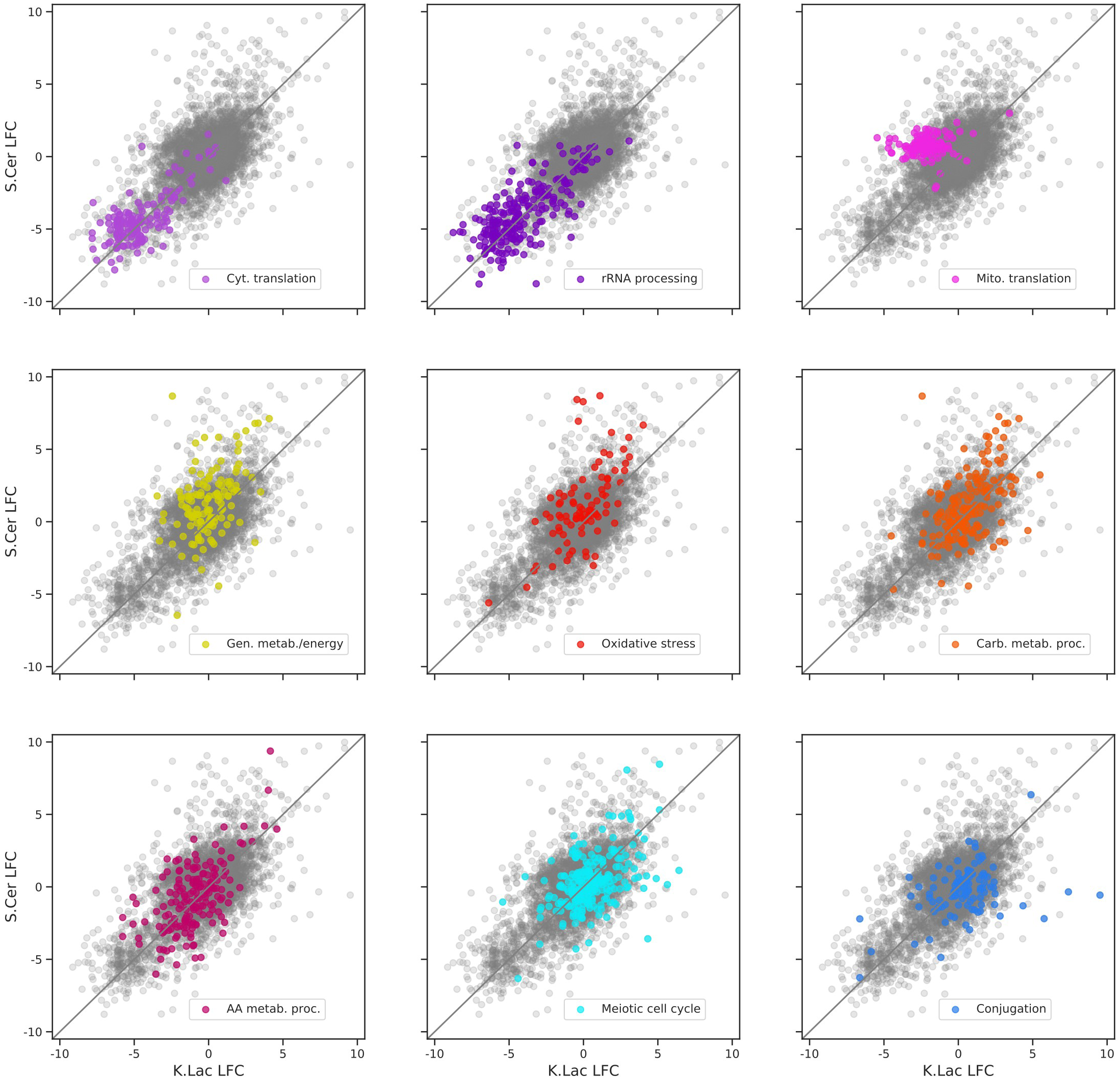
Distribution of LFC for selected GO-terms enriched in genes activated or repressed by PKA inhibition in *S. cerevisiae, K. lactis*, or both species. Log Fold Change (LFC) comparing RNA sequencing data collected from strains in which PKA-AS was inhibited with 3uM 1-NMPP1 versus DMSO controls. Data was collected after 50 min in both *S. cerevisiae* (y-axis) and *K. lactis* (x-axis) following administration of the drug. LFC values are only shown for genes that had orthologs in both species. Colored datapoints indicate all genes whose *S. cerevisiae* ortholog is a member of selected GO-Slim terms identified in Table S1. All other genes are colored grey.

**Figure S4:**
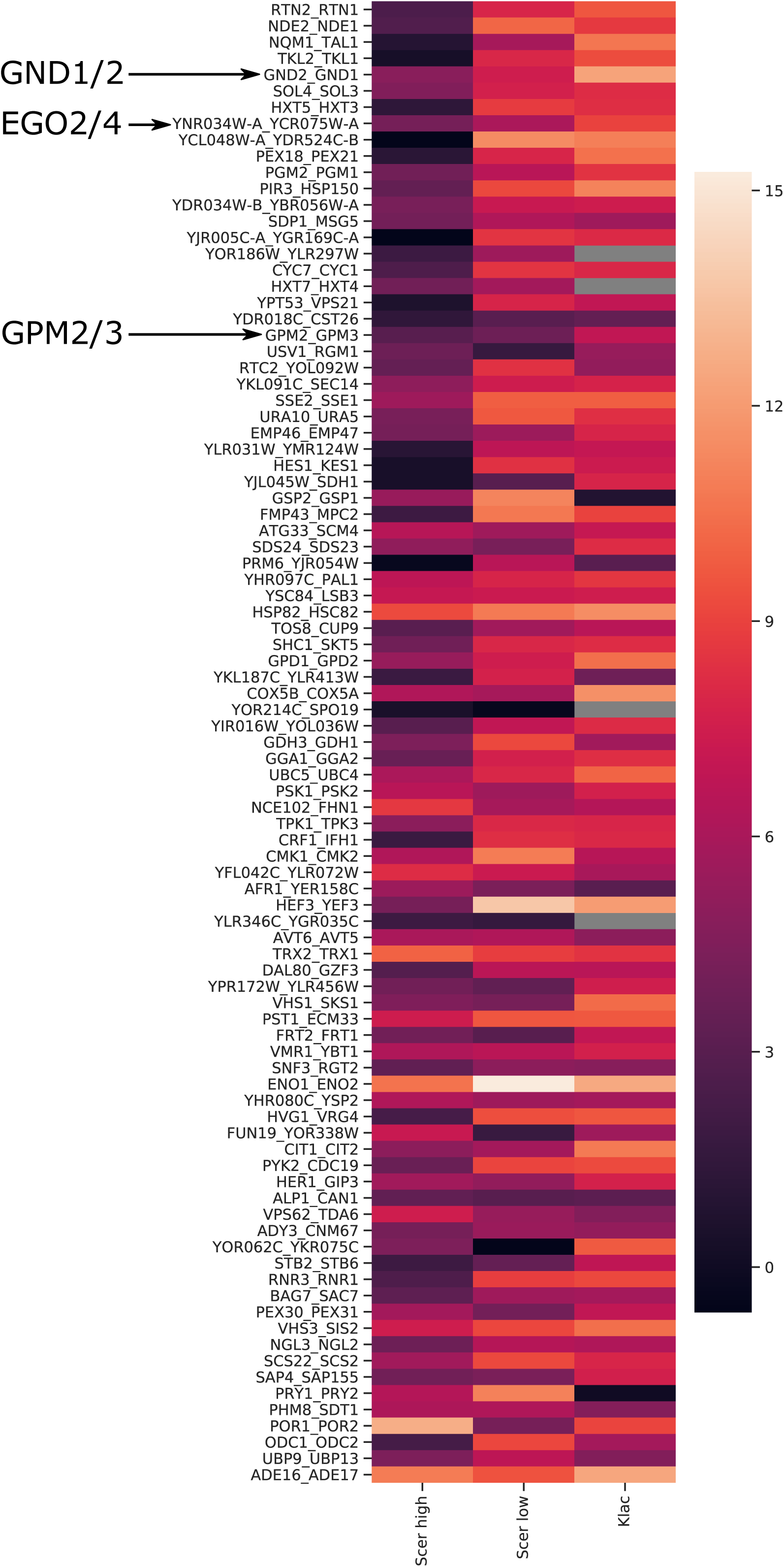
Basal expression (rlog) for DE*_PKA_* low and high LFC ohnologs and their shared orthologs in *K. lactis*. Average rlog data from PKA-AS strains with no drug during exponential growth are shown. Gray boxes indicate missing orthologs in *K. lactis*.

**Figure S5:**
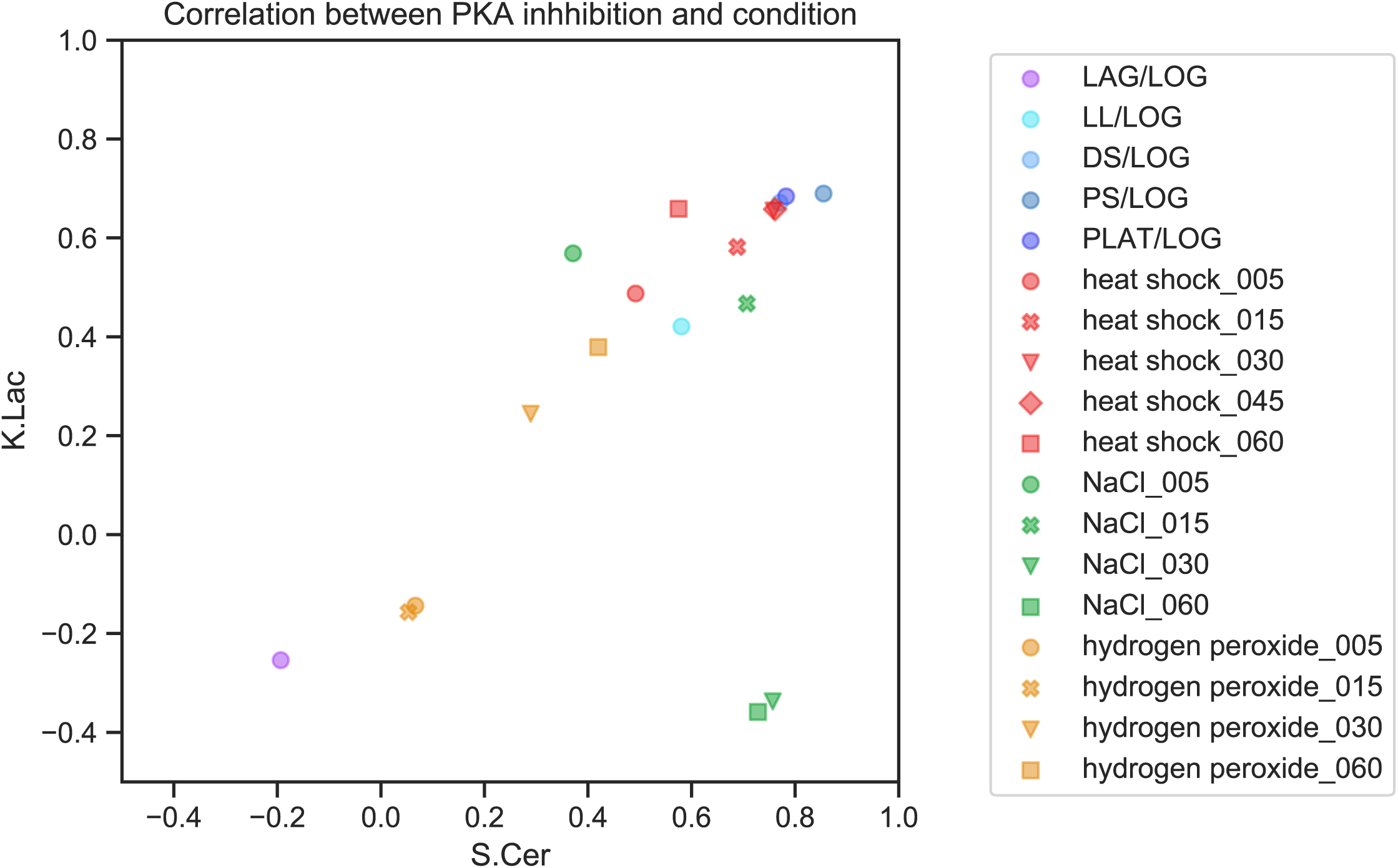
The stress conditions Diauxic Shift (DS), Post Diauxic Shift (PS), Plateau (PLAT), and heat shock at 30 and 45 minutes are most closely correlated with PKA inhibition in both *K. lactis* and *S. cerevisiae*. Pearson’s correlation coefficient between normalized LFC for the indicated condition from (Thompson et al., 2013) or (Roy et al., 2013) and our PKA inhibition data is plotted for *S.cerevisiae* (x-axis) and *K.Lactis* (y-axis).

**Figure S6:**
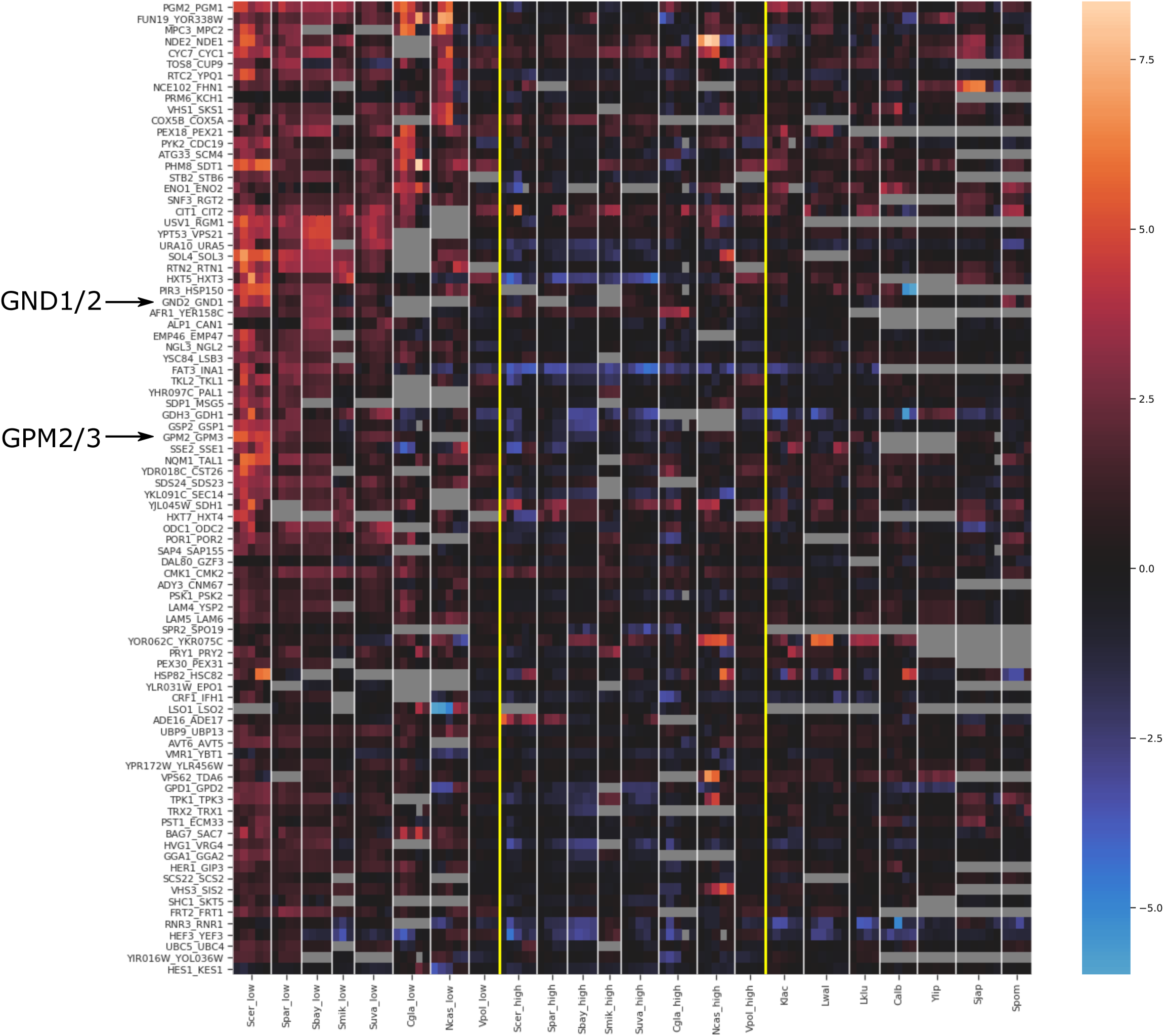
Induction of DE_PKA_ orthologs in response to stresses related to PKA inhibition. Normalized LFC values from the PKA-related stress conditions from Figure S5 are shown for the orthologs of each paralog pair in DE_PKA_. Yellow lines separate pre-WGH species (on the right) and post-WGH species (two groups on the left). Syntenic orthologs of hight-LFC ohnologs are on the left, and syntenic orthologs of low-LFC ohnologs are in the center. Where there are three columns, the conditions are ‘DS/LOG’, ‘PS/LOG’, and ‘PLAT/LOG’ from (Thompson et al., 2013) and where there are five bars, the conditions are those three conditions plus ‘heat shock_030’ and ‘heat shock_045’ from (Roy et al., 2013). *S. pombii* had the three growth conditions and ‘heat shock_30’. The rows are sorted such that the average LFC across conditions in the syntenic orthologs of the high-LFC ohnolog is more conserved towards the top and less conserved towards the bottom. The ohnolog pairs (YDR034W-B, YBR056W-A), (YCL048W-A, YDR524C-B), (CIS1, YGR035C), (EGO4, EGO2), and (YOR186W, YLR297W) in DE_PKA_ did not have sufficient data in the microarray experiments to be included in the analysis.

**Figure S7:**
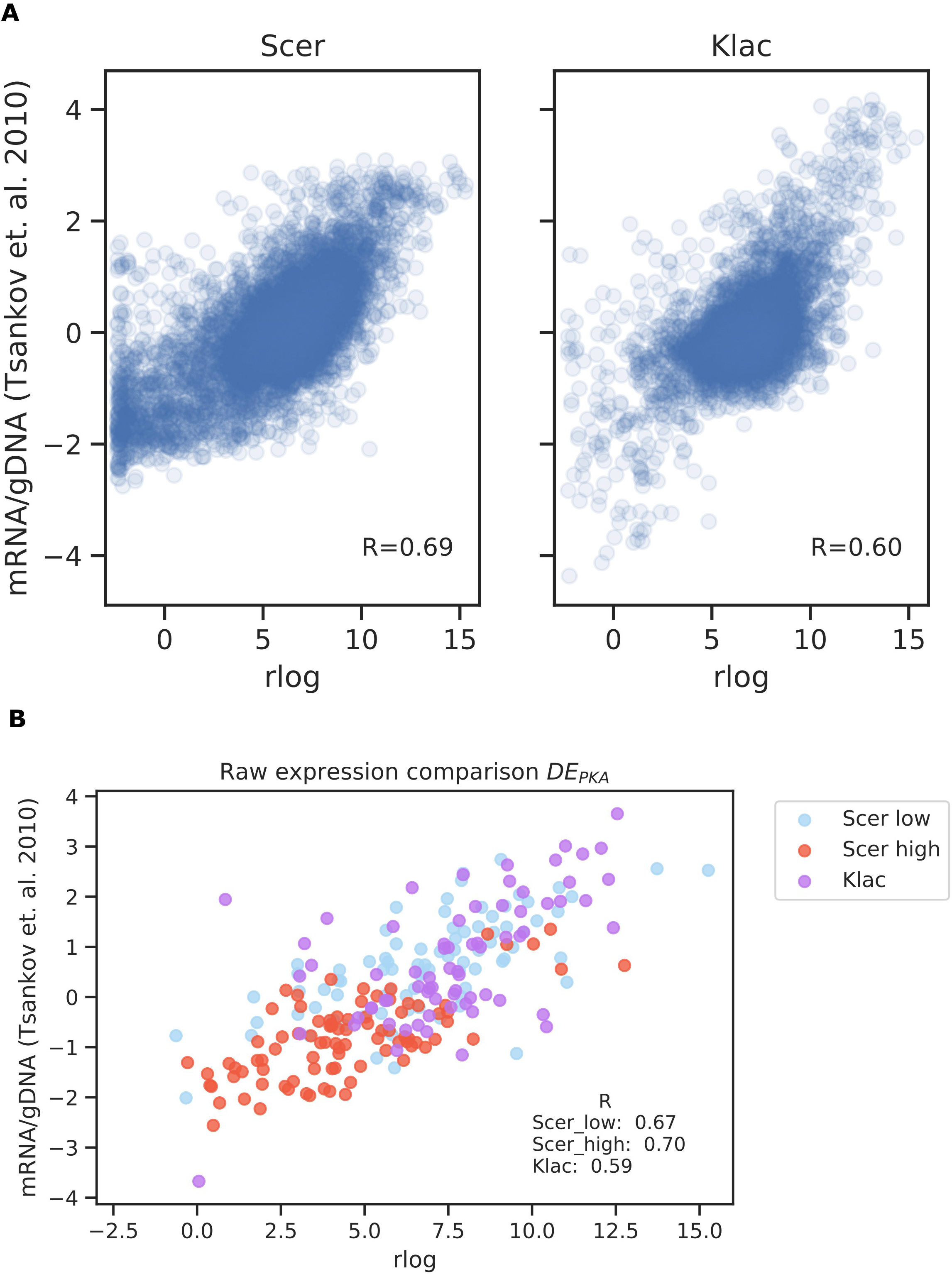
Basal expression from RNA-seq experiments is correlated with basal expression data from (Tsankov et al., 2010). (A) rlog data from a PKA-AS under exponential growth in YPD with no drug is shown on the x-axis, and normalized basal expression from (Tsankov et al., 2010) (See Materials and Methods for details on normalization) is shown on the y-axis for all genes in *S. cerevisiae* (left panel) and *K. lactis* (right panel). (B) The same data as in (A) for Low-LFC (blue) and High-LFC (red) DE_PKA_ ohnologs from *S. cerevisiae* and their shared *K. lactis* orthologs (purple) are shown.

**Figure S8:**
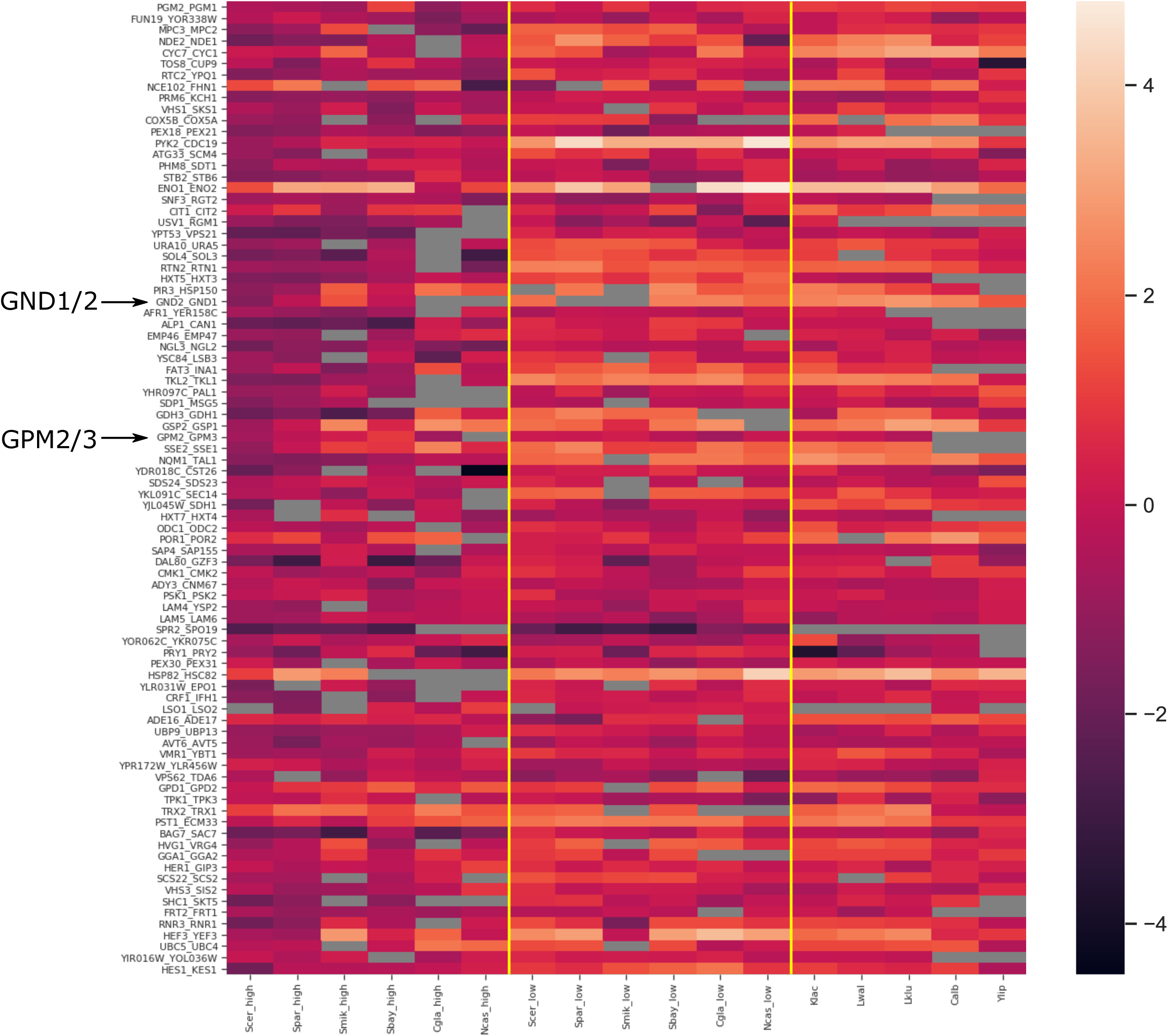
Basal expression of DE_PKA_ orthologs for 11 budding yeast species. Normalized expression data (see Materials and Methods) for each species is shown from (Tsankov et al., 2010). Rows and columns are ordered as in Fig S6.

**Figure S9:**
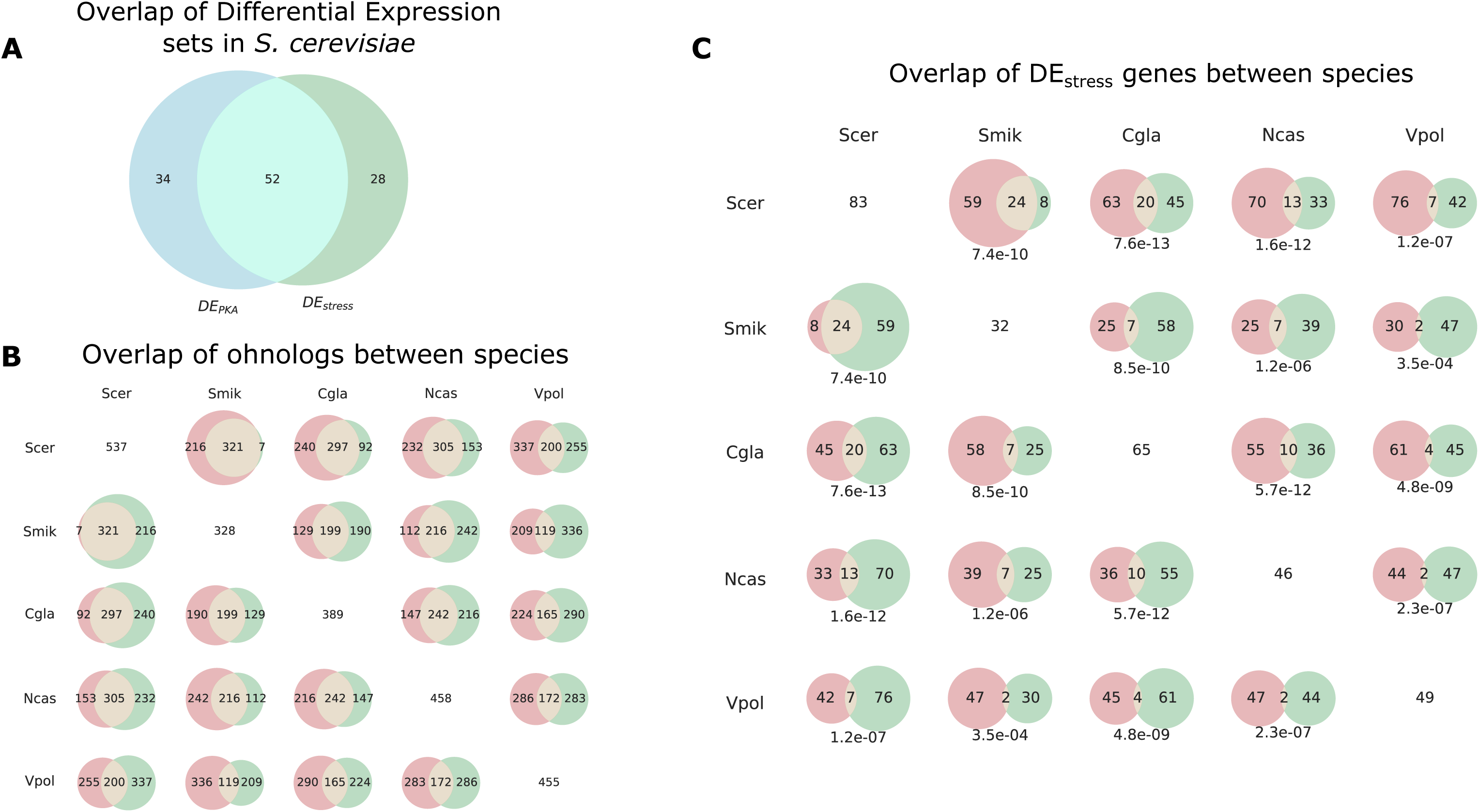
Non-overlapping sets of genes are differentially expressed in response to stress in different post-WGH species. We can define ohnolog pairs that have one member activated by PKA-related stress conditions (Fig S5) and which have differential expression in a similar manner as we defined DE_PKA_, which we denote for a particular species as 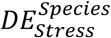 (See Materials and Methods). (A) Overlap between *DE_PKA_* ohnolog pairs (which are defined in *S. cerevisiae)* and 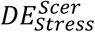. *DE_Stress_* ohnolog pairs were defined in each species based on the average normalized LFC across the 3-5 PKA inhibition related conditions (LFC_est_). We first identified all ohnolog pairs in which one ohnolog was activated (LFC_est_>1.8), and the other was not (LFC_est_ < 1.6). We retained ohnolog pairs in which the difference in LFC_est_ between the activated and non-activated ohnolog was greater than 1.8. The overlap between *DE_PKA_* and 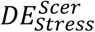 (52 shared ortholog pairs) is significantly more than would be expected by chance (p=1.29E-27, Fisher’s exact test) (B) Overlap of all ohnologs that have data in (Roy et al., 2013; Thompson et al., 2013) between species as determined from YGOB pillars (Byrne and Wolfe, 2005). The number on the diagonal is the total number of ohnologs with data in that species. (C) Overlap between *DE_Stress_* sets for various pairs of species. Shown below each Venn diagram is the p-value for a Fisher’s exact test on the null hypothesis that this overlap is expected based on the overlap of all ohnologs with data (shown in (B)) for any two given species. The number on the diagonal is the total number of ohnolog pairs in *DE_Stress_* for that species. There is little overlap between orthologs of the *DE_Stress_* sets defined for different species. This is partially a result of the fact that the set of ohnolog pairs that is retained decreases as the evolutionary distance between species increases for post-WGH species (Scannell et al., 2007). However, the overlap of differentially expressed ohnolog pairs is even smaller than would be expected given the total percentage of retained ohnologs between species. Thus, there is no reason to expect a priori that the same conservation patterns for LFC and basal expression would hold for differentially expressed ohnolog pairs defined by their expression in distantly related post-WGH species 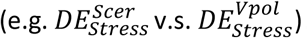. Species abbreviations: Scer = *Saccharomyces cerevisiae*, Spar = *Saccharomyces paradoxus*, Sbay = *Saccharomyces bayanus,* Smik = *Saccharomyces mikatae*, Suva = *Saccharomyces uvarum*, Cgla = *Candida glabrata*, Ncas = *Nakaseomyces castellii,* Vpol = *Vanderwaltozyma polyspora*.

**Figure S10:**
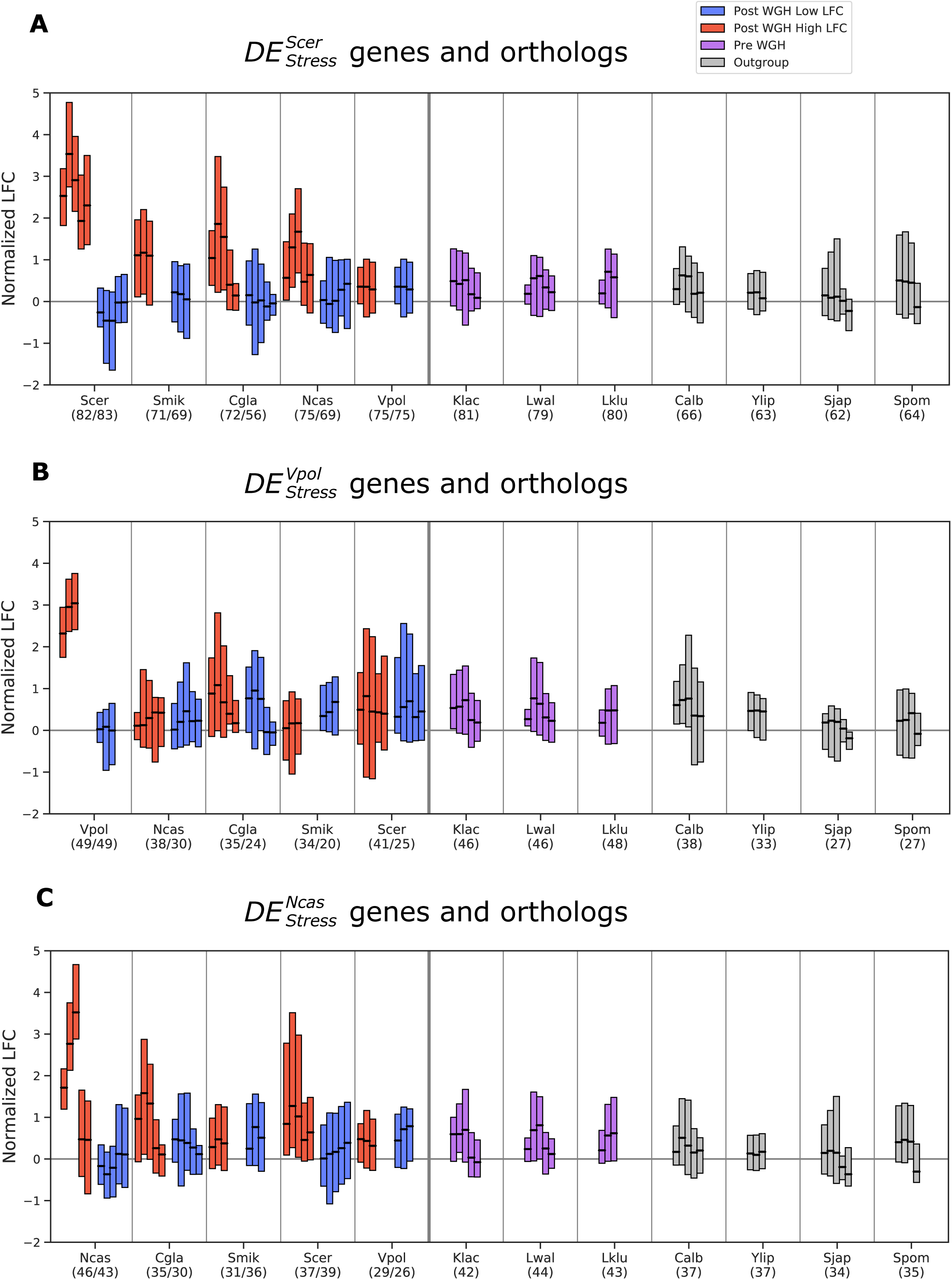
The conclusion that PKA induction is the derived phenotype is independent of the species in which *DE_Stress_* ohnolog pairs are defined. Distribution of LFC values for PKA related stress conditions (as in Fig 3B) is shown, except focusing on 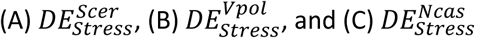. Boxplots show median and Q1-Q3 range for normalized LFC (see Materials and Methods) for gene expression data from (Roy et al., 2013; Thompson et al., 2013) for the indicated species for the stress conditions most correlated to PKA inhibition in *S. cerevisiae* and *K. lactis* from (Fig S5). Where there are three bars, the conditions are ‘DS/LOG’, ‘PS/LOG’, and ‘PLAT/LOG’ from (Thompson et al., 2013) and where there are five bars, the conditions are those three conditions plus ‘heat shock_030’ and ‘heat shock_045’ from (Roy et al., 2013). *S. pombii* did not have ‘heat shock_45’. Boxplots for data from *DE_Stress_* genes from (A) *S. cerevisiae*, (B) *V. polymorpha* and (C) *N. castelliii* are shown towards the left side of each panel and boxplots for data from the orthologs in indicated species of those *DE_Stress_* genes are shown to the right. Blue and red indicate low-LFC and high-LFC ohnologs (respectively) and their syntenic orthologs in post-WGH species. Purple and grey bars are for the shared orthologs in Pre-WGH *Saccharomycetaceae* species and outgroups respectively. Numbers in parentheses indicate the number of retained orthologs. Syntenic ortholog assignment for post-WGH species is based on the YGOB database (Byrne and Wolfe, 2005).

**Figure S11:**
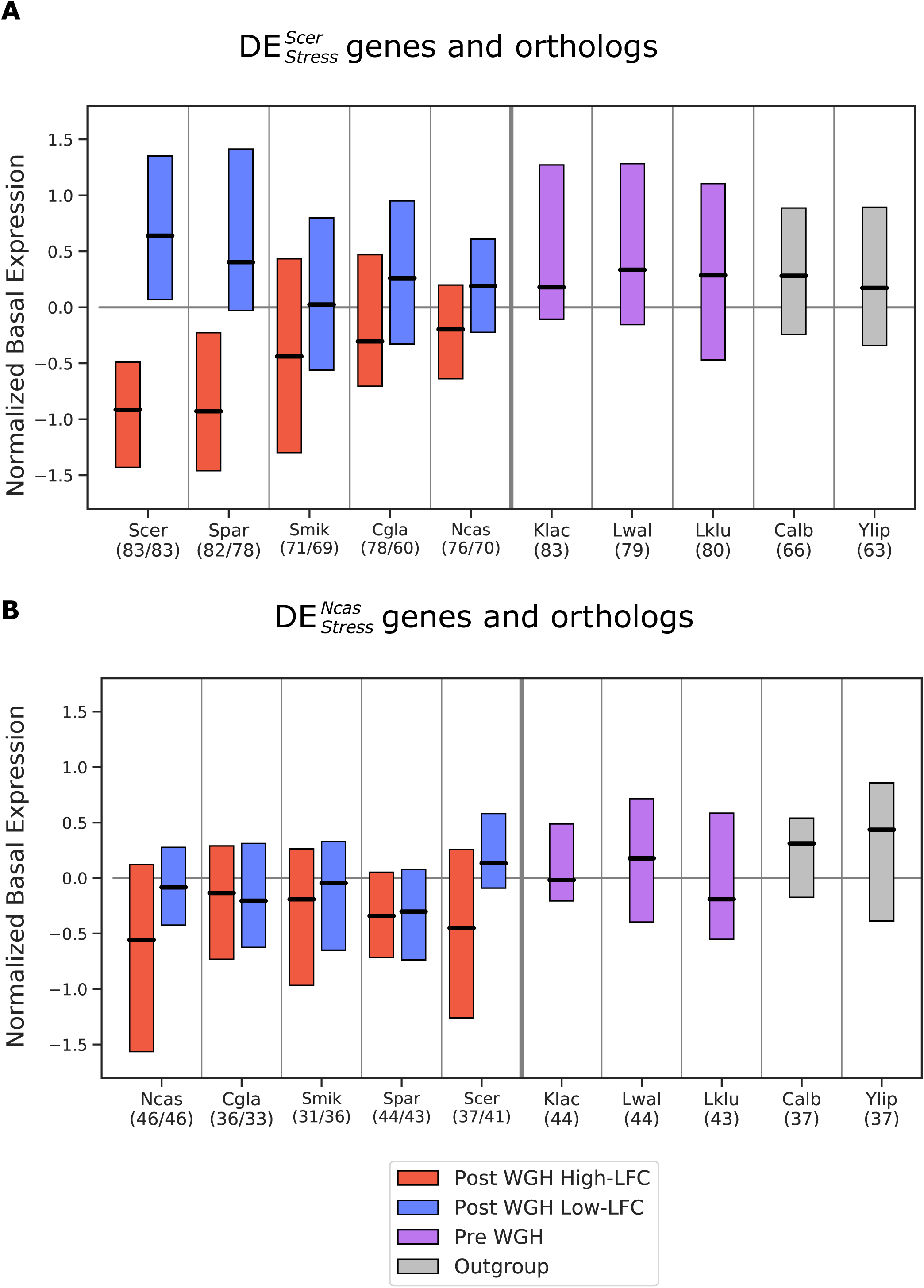
The conclusion that high basal expression is the ancestral phenotype is independent of the species in which *DE_Stress_* ohnolog pairs are defined. Distribution of basal expression values as in Fig 3C is shown, except focusing on 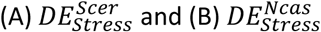. Boxplots showing median and Q1-Q3 range of normalized raw expression data (see Materials and Methods) are shown from microarray experiments comparing mRNA under exponential growth conditions to genomic DNA from (Tsankov et al., 2010). Boxplots for data from *DE_Stress_* genes from (A) *S. cerevisiae* and (B) *N. castelliii* are shown towards the left side of each panel and boxplots for data from the orthologs in indicated species of those *DE_Stress_* genes are shown to the right. Blue and red indicate low-LFC and high-LFC ohnologs (respectively) and their syntenic orthologs in post-WGH species. Purple and grey bars are for the shared orthologs in pre-WGH *Saccharomycetaceae* species and outgroups respectively. Syntenic ortholog assignment for post-WGH species is based on the YGOB database (Byrne and Wolfe, 2005).

**Figure S12:**
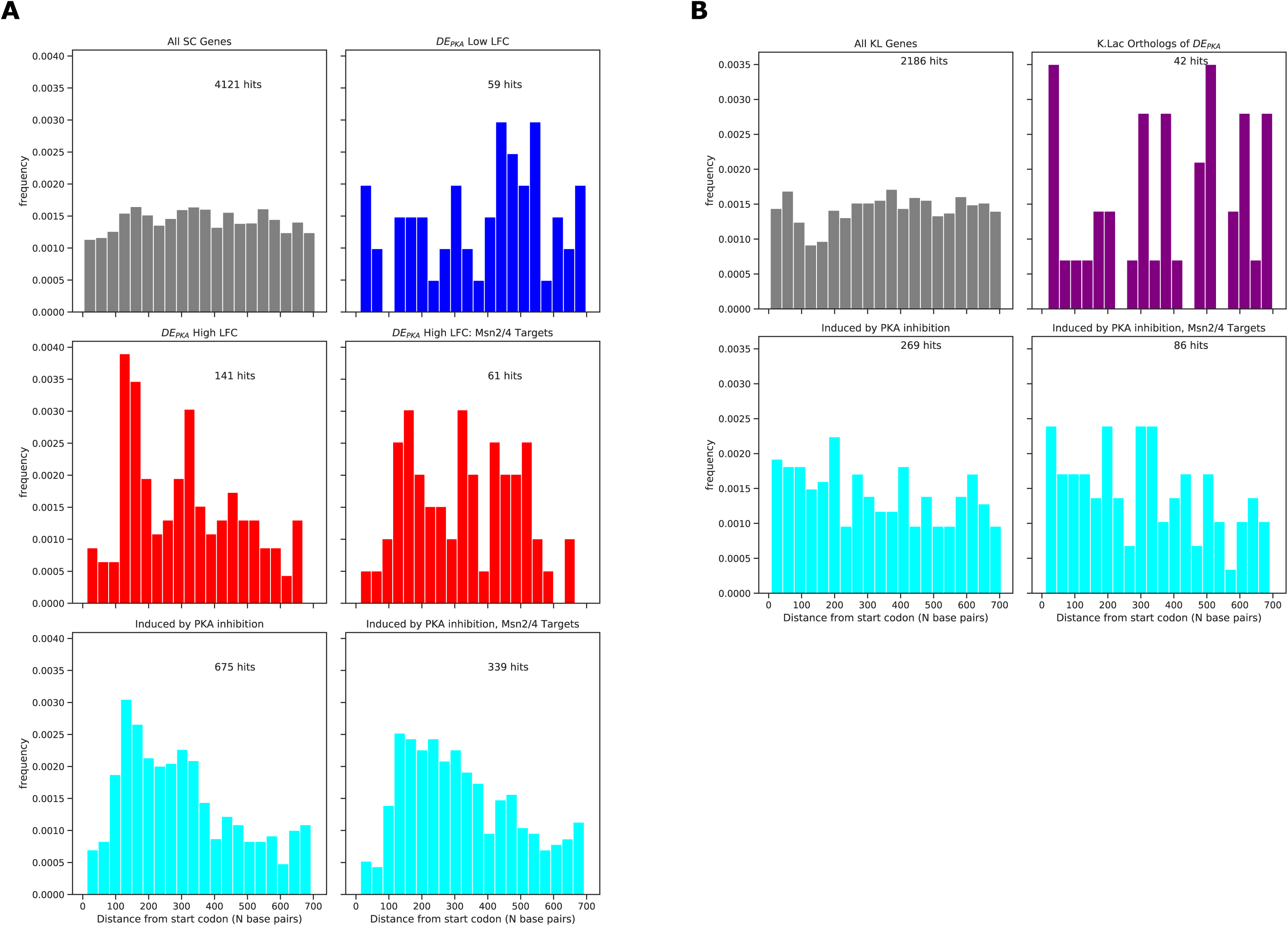
The STRE is localized closer to the start codon in the promoters of targets of PKA inhibition in *S. cerevisiae*. Distribution of STRE distance from start codon for indicated gene sets for (A) *S. cerevisiae* and (B) *K. lactis*.

**Figure S13:**
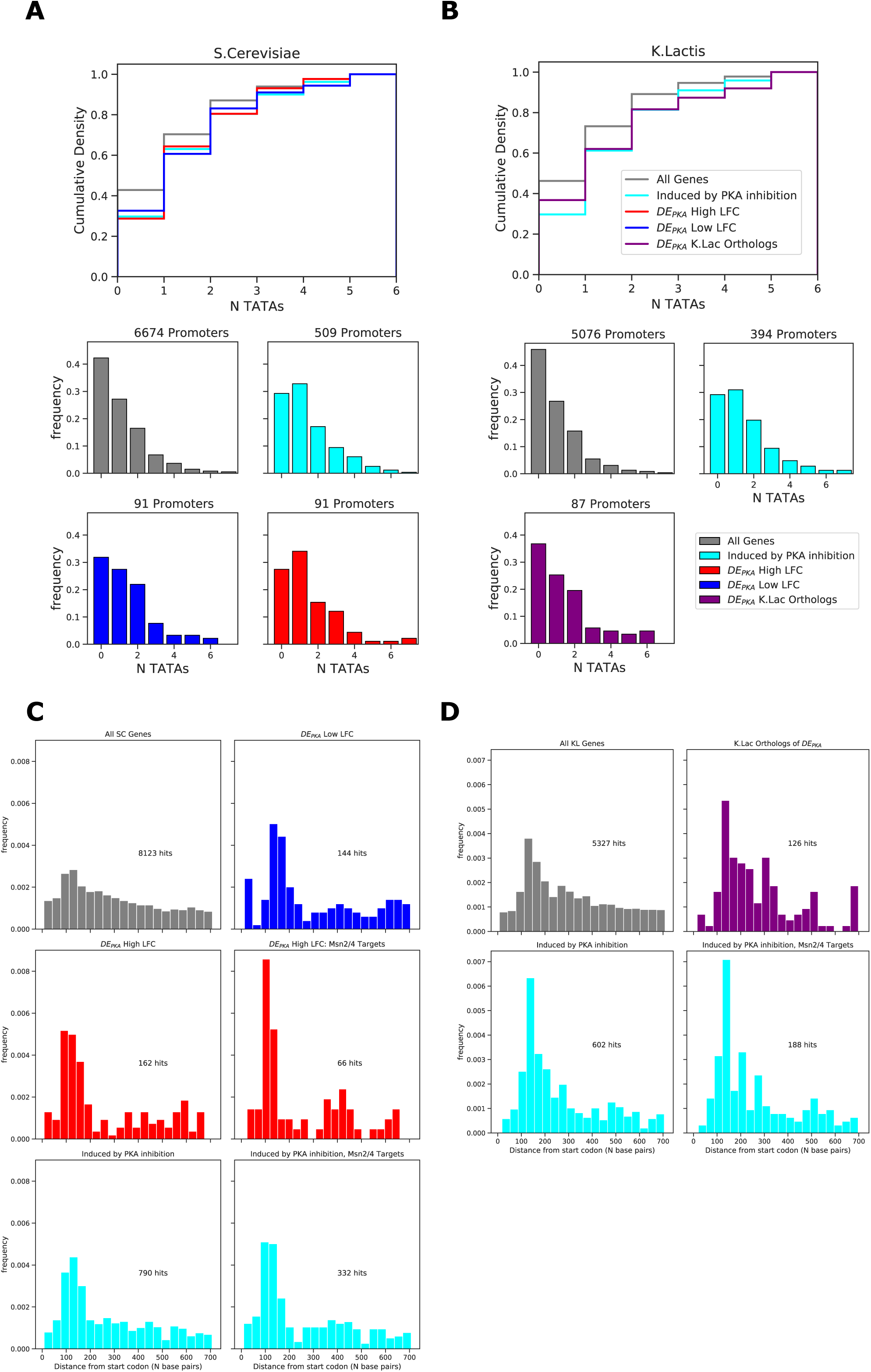
The TATA box is enriched in the promoters of targets of PKA inhibition, as well as in orthologs of DE_PKA_ genes in *S. cerevisiae* and *K. lactis*. (A) Distribution of number of TATA boxes in the promoters of indicated sets in *S. cerevisiae*. (B) same for *K. lactis*. (C) Distribution of the distance of TATA boxes from the start codon in the promoters of indicated sets for *S. cerevisiae* and (D) *K. lactis*.

**Figure S14:**
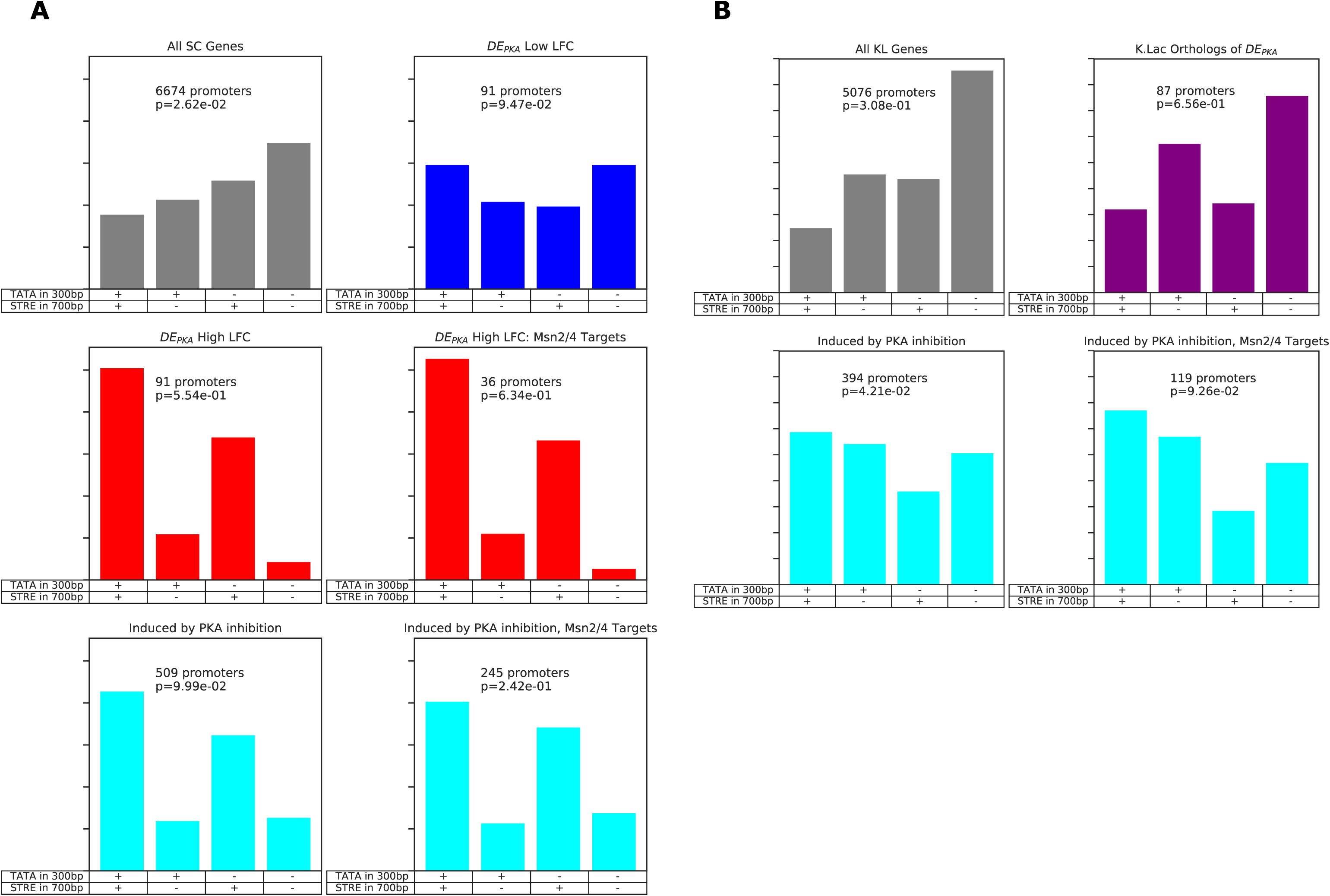
There is a high percentage of promoters with both TATA boxes and STREs in genes induced by PKA in *S. cerevisiae*, which is expected based on the enrichment for both motifs in that set. Percentages of promoters in the indicated sets with one or more STRE in combination with one or more TATA boxes in the 300bases upstream of the start codon for (A) *S. cerevisiae* and (B) *K. lactis*.

**Figure S15:**
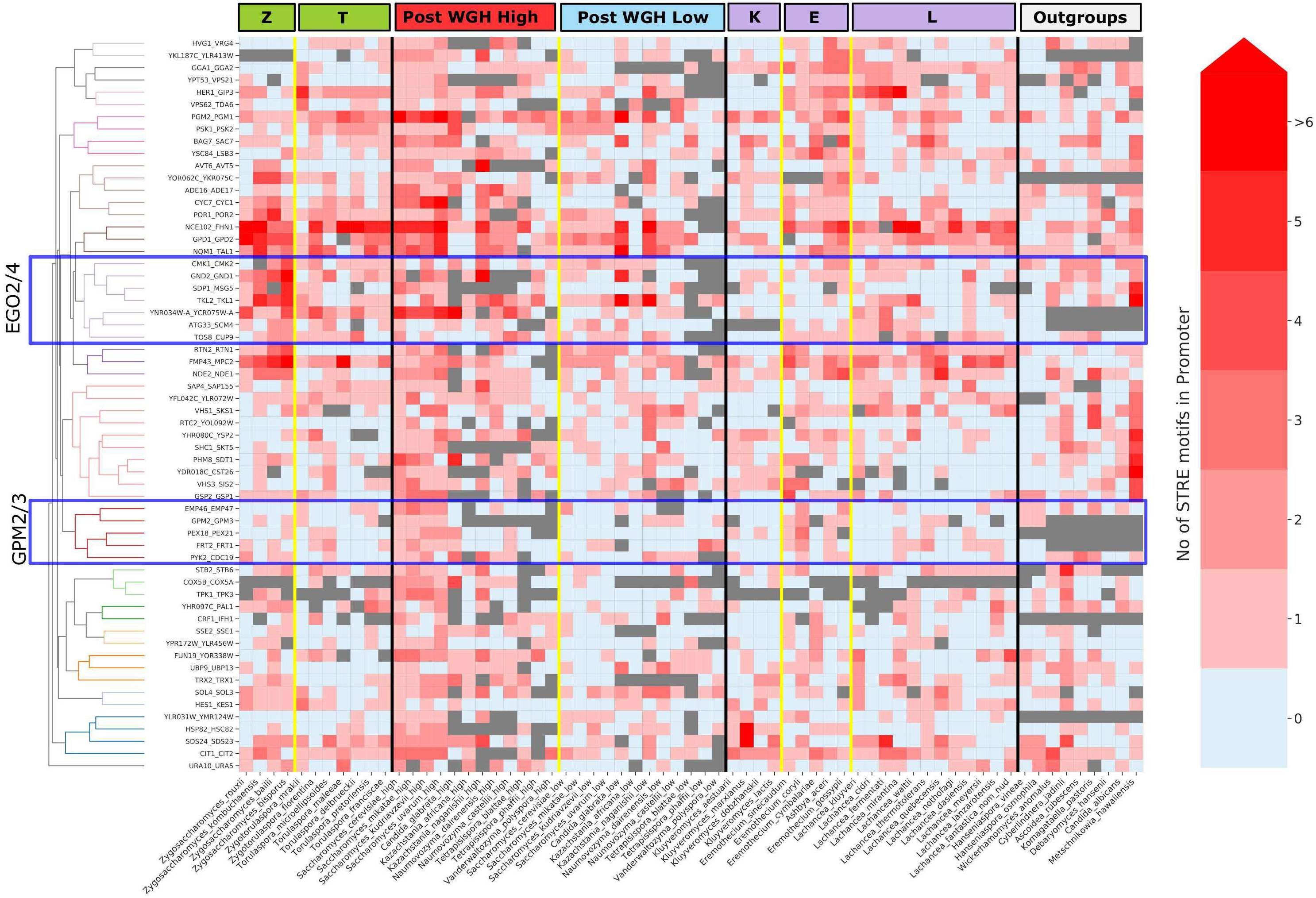
STRE counts in the promoters of orthologs of select DE_PKA_ genes. The number of STREs in the promoters of the orthologs of the subset of DE_PKA_ genes considered for Fig 5 are shown. Grey indicates either no ortholog exists or no promoter was found in the dataset. Columns are different species and rows are clustered based on the number of STREs in pre-WGH species (ZT branch, KLE branch, and outgroups). The dendrogram for the hierarchical clustering is shown to the left. Clusters highlighted in Fig 6B are indicated in blue boxes.

**Figure S16:**
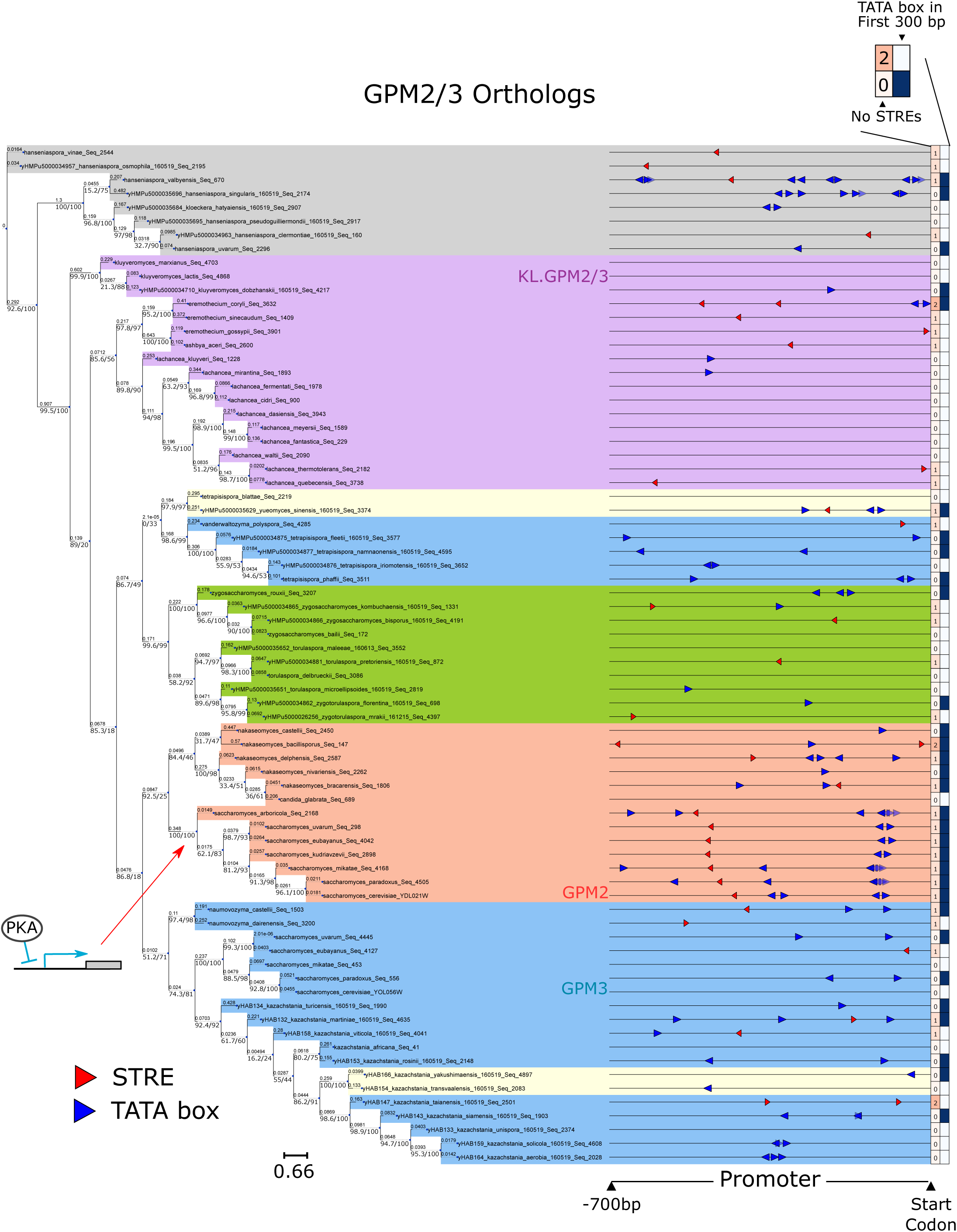
GPM2/3 are an example of a differentially induced pair of ohnologs in which the STRE arose in the promoter of the DE_PKA_ high-LFC ohnolog following the WGH. Phylogenetic tree of all orthologs of GPM2/3 from the *Saccharomycetaceae* clade from (Shen et al., 2018) plotted alongside each gene’s promoter (700 bp upstream of the start codon) with STRE (red triangle) and TATA box (blue triangles) motifs highlighted. The arrow indicates the putative point at which the STRE was gained. The first column of boxes after each promoter represents the number of STREs and the second columns of boxes indicates whether there is a TATA box within 300 bases of the start codon. Phylogeny is determined from a multiple sequence alignment of the protein sequences (see Materials and Methods). Support values (bootstrap/alrt) are shown to the left and below each branch point, and branch lengths (amino acid substitutions/site) are shown above. Shading represents different groups of species; blue = Post-WGH, syntenic ortholog to low-LFC ohnolog; red = Post-WGH, syntenic orthologs to high-LFC ohnolog; yellow=Post-WGH, synteny not determined; green= ZT; light purple=KLE; dark purple=other Pre-WGH; grey=outgroups.

**Figure S17:**
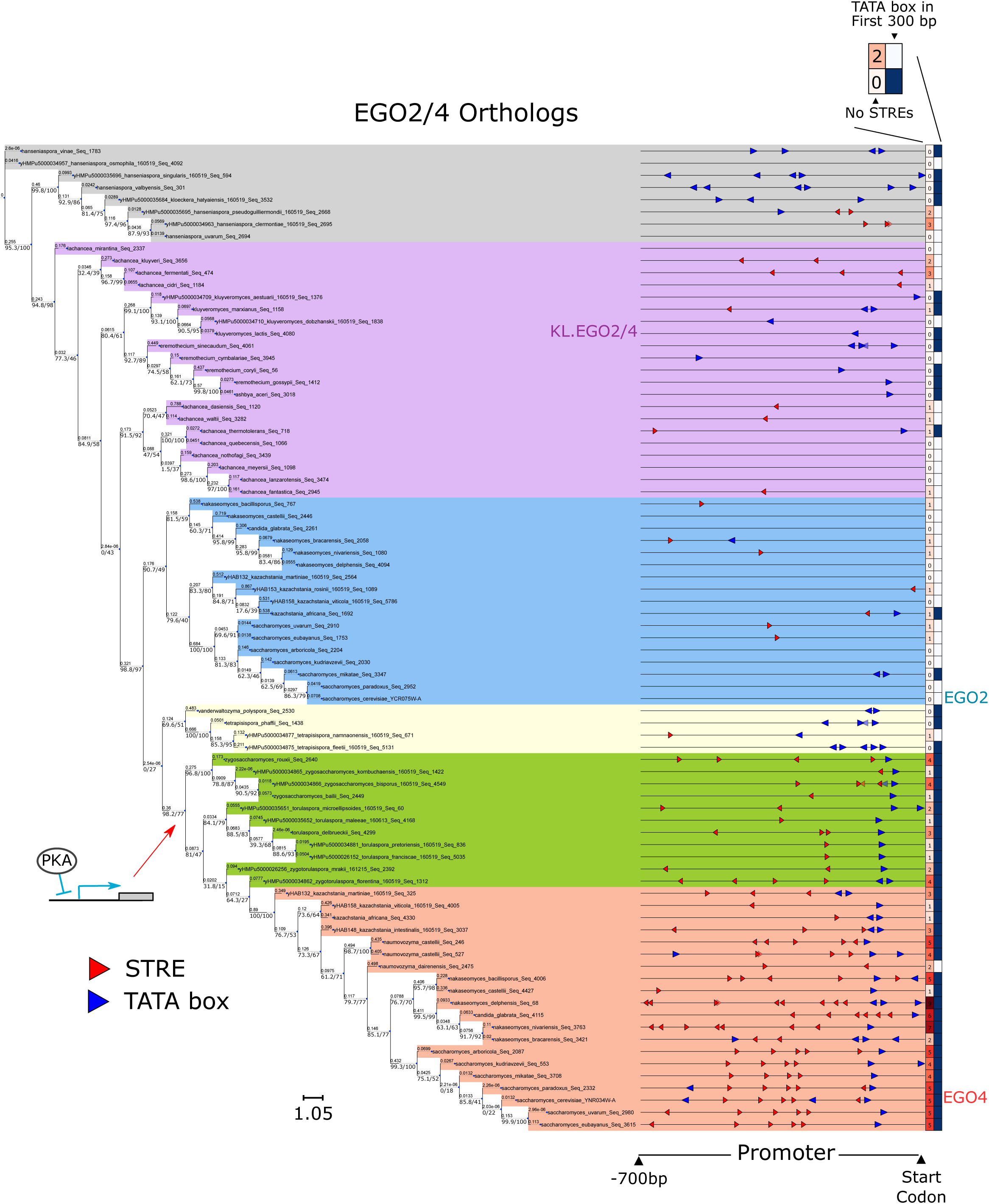
EGO2/4 are examples of a differentially induced ohnolog pair in which the STRE arose in the ZT branch prior to the WGH. Phylogenetic tree of all orthologs of EGO2/4 from the *Saccharomycetacea*e clade from (Shen et al., 2018) plotted alongside each gene’s promoter. Conventions and nomenclature are as in Fig S16.

**Figure S18:**
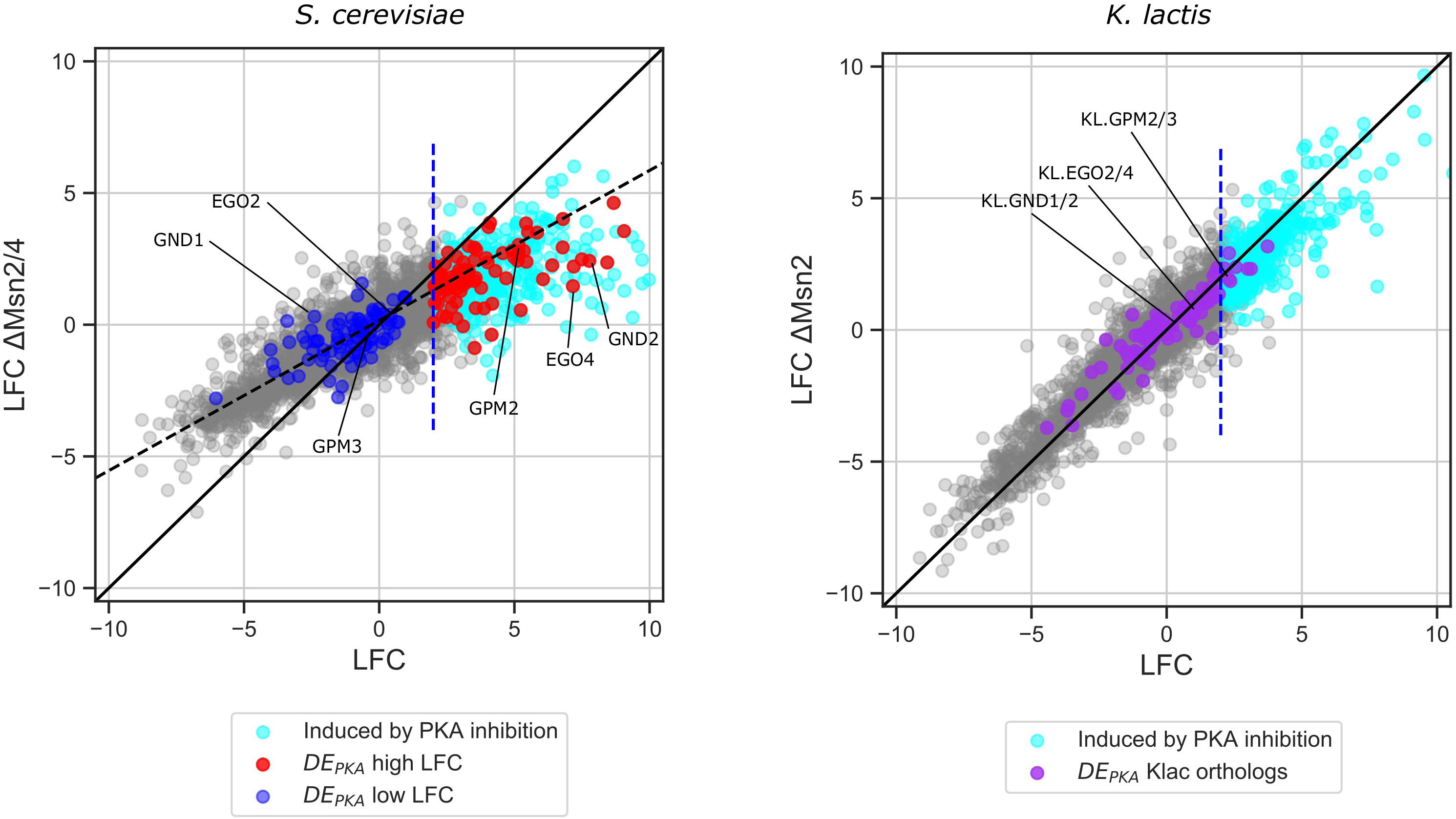
The expression of many DE_PKA_ high LFC ohnologs depends on Msn2/4. RNA seq data for PKA-AS strains with and without (A) Msn2/4 in *S. cerevisiae* or (B) their shared ortholog in *K. Lactis* in the presence of 3µM 1-NM-PP1 at 50 minutes. The solid black line is 1:1. The dashed line in (A) is a regression line based on the genes (slope 0.57, intercept 0.16) with a negative LFC in WT cells to illustrate a general decrease in the response to PKA for both repressed and activated genes in ΔMsn2/4 cells.

**Figure S19:**
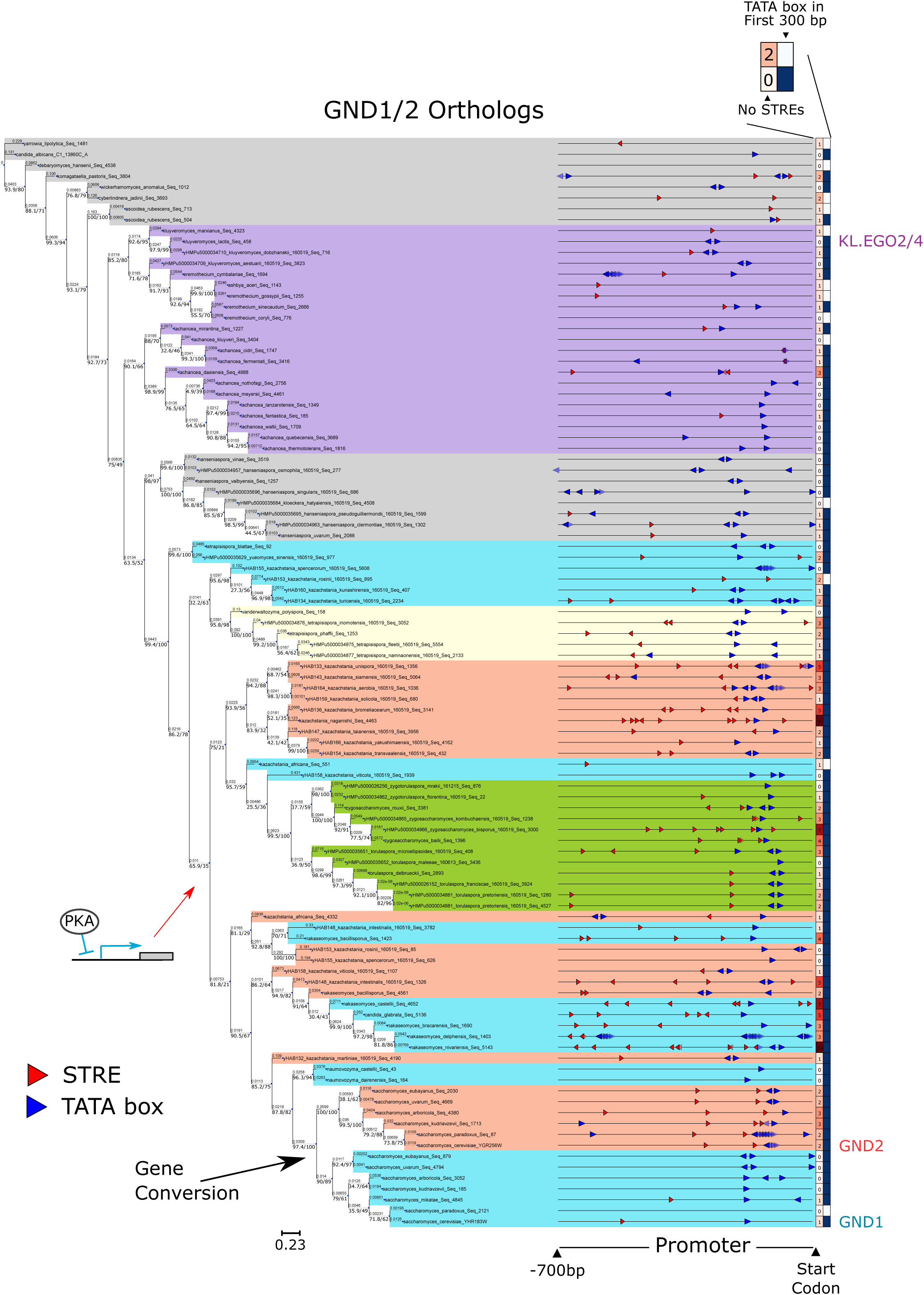
GND1/2 are examples of a differentially induced ohnolog pair in which there was a gene conversion for the protein, but in which the STRE appears to have arisen in the ZT branch prior to the WGH. Phylogenetic tree of all orthologs of GND1/2 from the *Saccharomycetaceae* clade from (Shen et al., 2018) plotted alongside each gene’s promoter. Conventions and nomenclature are as in Fig S16.

